# Cerebellar Control of a Unitary Head Direction Sense

**DOI:** 10.1101/2021.07.08.451624

**Authors:** Mehdi Fallahnezhad, Julia Le Méro, Xhensjana Zenelaj, Jean Vincent, Christelle Rochefort, Laure Rondi-Reig

## Abstract

Head direction (HD) cells, key neuronal elements in the mammalian’s navigation system, are hypothesized to act as a continuous attractor network, in which temporal coordination between cell members is maintained under different brain states or external sensory conditions, resembling a unitary neural representation of direction. Whether and how multiple identified HD signals in anatomically separate HD cell structures are part of a single and unique attractor network is currently unknown. By manipulating the cerebellum, we identified pairs of thalamic and retrosplenial HD cells that lose their temporal coordination in the absence of external sensory drive, while the neuronal coordination within each of these brain regions remained intact. Further, we show that distinct cerebellar mechanisms are involved in the stability of direction representation depending on external sensory conditions. These results put forward a new role for the cerebellum in mediating stable and coordinated HD neuronal activity toward a unitary thalamocortical representation of direction.

## INTRODUCTION

The sense of direction in the mammalian brain is thought to be encoded by the activity of HD cells, neurons that robustly increase their firing when the animal’s head points in a particular direction (Ranck, 1984; Taube et al., 1990a). Population activity of such neurons is postulated to perform as a continuous attractor network in which temporal coordination between HD cells is preserved independent of brain state or sensory conditions, creating an internally-organized neural compass for direction representation (Park et al., 2019; Peyrache et al., 2015; Seelig and Jayaraman, 2015). For example, HD cells exhibit a preserved pairwise temporal coordination during wake and sleep within and between anatomically separate HD cell structures (Peyrache et al., 2015), during mismatch external sensory conditions (Park et al., 2019; Yoganarasimha et al., 2006), before anchoring to the external world (Bassett et al., 2018; Bjerknes et al., 2015), when the HD system is decoupled from the vestibular system (Muir et al., 2009; Yoder and Taube, 2009) or from its potential velocity inputs (Butler et al., 2017). The existence of such a continuous HD attractor network is more directly shown in insects, in which an artificially-induced activity within a subsample of neurons representing heading signal leads into a rapid shift of activity bump centered in the stimulation site (Kim et al., 2017). The findings in insects and rodents altogether suggest that the mammalian HD system may perform as a continuous attractor network (Knierim and Zhang, 2012; Mcnaughton et al., 1996, 1991; Redish et al., 1996; Samsonovich and McNaughton, 1997; Skaggs et al., 1995) and potentially resembles a single, and unique internal compass (Green et al., 2017; Hulse and Jayaraman, 2020; Kim et al., 2017; Park et al., 2019). Conversely, some recent studies suggest the potential presence of multiple heading signals in the mammalian brain, such as local- and global-cue anchoring HD cells in the retrosplenial cortex (Jacob et al., 2016), theta versus non-theta HD cells with different anchoring responses to the visual-cue manipulations in the medial entorhinal cortex (Kornienko et al., 2018) as well as the presence of a vestibular-independent heading signal in the hippocampus (Acharya et al., 2016). However, an incoherent change in the preferred firing direction (PFD) of a subsample of HD cells or a varied directional firing pattern within HD cell assemblies in response to a mismatch sensory condition does not necessarily imply the presence of multiple attractor networks. Indeed, under a conflict between the local and global cue, HD cells start to anchor differently to either of these cues (Knight et al., 2014; Park et al., 2019), while the sub-second temporal correlation structure between all HD cell pairs is preserved (Park et al., 2019). These results indicate that temporal coordination between HD cell assemblies remains intact despite varied allocentric directional responses. Therefore, whether and how HD signals across separate brain structures remain part of a single and unique continuous attractor network independent of any external sensory drive is currently unknown.

To function as a neural compass, the internally organized HD cell assemblies is anchored to the external world by utilizing multimodal sensory information. Specifically, both vestibular and motor inputs are shown to be essential for stable allocentric HD representation (Muir et al., 2009; Stackman, 2003; Stackman and Taube, 1997; Stackman et al., 2002; Valerio and Taube, 2016; Yoder and Taube, 2009; Zugaro et al., 2001). Landmark information (e.g., visual, olfactory, auditory) are also used for updating the HD representation (Clark et al., 2009; Goodridge and Taube, 1995; Goodridge et al., 1998). It is accordingly clear that multimodal sensory integration is critical for stable HD cell activity patterns (Cullen and Taube, 2017; Taube, 2007; Taube and Bassett, 2003; Yoder et al., 2011). However, it is not known how and where in the brain the multimodal information necessary for the stable representation of direction is processed, filtered, and fed to the HD circuitry.

Cerebellum has long been considered in processing sensory information (Gao et al., 1996; Ishikawa et al., 2015; Snider and Stowell, 1944), sensory adaptation, and sensory prediction error (Bastian, 2006; Diedrichsen et al., 2005; Imamizu et al., 2000; McDougle et al., 2016; Pasalar et al., 2006; Popa and Ebner, 2019; Shadmehr et al., 2010; Streng et al., 2018; Wolpert et al., 1998). The cerebellum also processes head-body orientations in a gravity reference frame (Angelaki et al., 2004; Dakin et al., 2018; Dugué et al., 2017; Laurens et al., 2013b, 2013a; Mackrous et al., 2019). However, there is no direct evidence showing whether and how this structure may influence HD cell firing and HD cell assemblies. We hypothesized that the cerebellum might play a critical role in processing the information required for stable HD cell activity patterns. Also, since the cerebellum is involved in temporal synchronization between distant structures (Lindeman et al., 2021; Liu et al., 2020; McAfee et al., 2019; Popa et al., 2013), we sought to examine the influence of the cerebellum on the coordination between different HD structures.

To this end, we employed two different cerebellar-specific transgenic mouse models, one with a deficiency in protein kinase C (PKC)-mediated long-term depression (L7-PKCI) and the other with a defect in protein phosphatase 2B (PP2B)-mediated potentiation (L7-PP2B), both specifically targeting Purkinje cell plasticity. Monitoring the simultaneous activity of HD cells from the retrosplenial cortex (RSC) and anterodorsal thalamic nucleus (ADN), we investigated the dynamics of single and ensemble HD cell activities both within and between cortical and subcortical HD cell networks under navigations by landmarks or idiothetic cues.

Here, we identify a novel role for the cerebellum in maintaining a single and stable direction representation across thalamocortical HD cell networks. We demonstrate the presence of two separate and potentially self-organized HD networks within the RSC and ADN structures, whose generic temporal coordination toward building a unitary representation of direction requires cerebellar computation. Further, we show that while the cerebellar PP2B-dependent mechanism is involved in HD cell anchoring to allocentric cues, cerebellar PKC-dependent processing is required for driving HD cells by idiothetic cues.

## RESULTS

### HD Cells are Unstable in L7-PKCI Mice When Relying on Self-Motion Cues

We performed extracellular single-unit recordings from HD cells in ADN and RSC of six L7-PKCI mice and seven control littermates (**Figure 1A**, **Table S1, and Figures S1 and S2**). Recordings were made while animals performed a sequence of six 10-min foraging sessions under different sensory conditions with 5-min inter-session resting intervals (except before the dark session - **Figure 1B**). The sessions included, in the following order, two standard light sessions in the presence of a proximal cue card (sessions L_1_ and L_2_), a 90° counterclockwise cue rotation session under light (rotation session R_3_), another standard light session (session L_4_) followed by a dark session with cue removal (no light no cue session D_5_), and finally a standard light session (session L_6_). Recorded spikes were sorted offline into single units, and the units resembling HD cell activity were identified either using a permutation test or by modeling the spiking in a generalized linear model framework (**Supplemental Experimental Procedures**).

**Figure 1.**
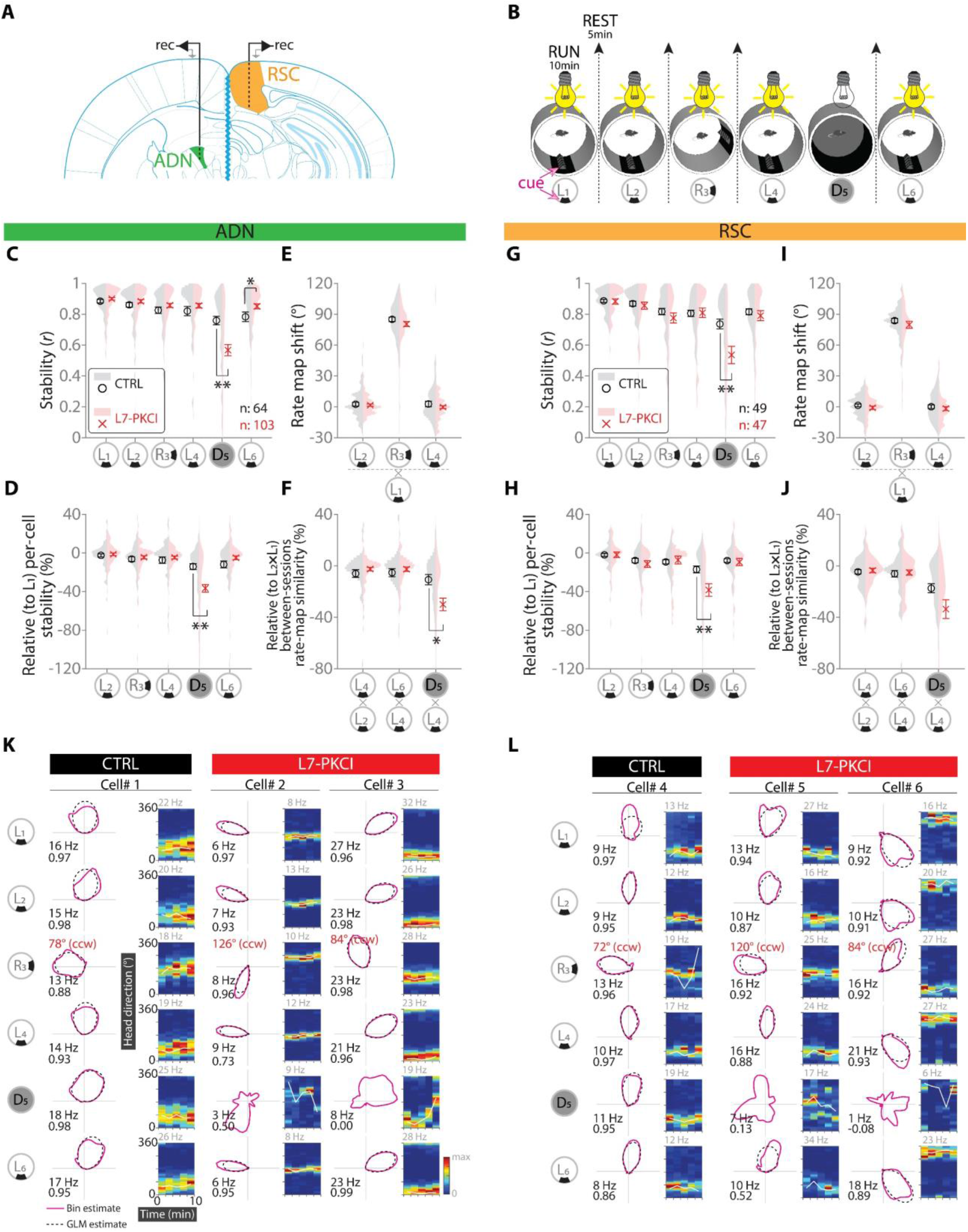
HD Cells are Unstable in L7-PKCI Mice When Relying on Self-Motion Cues. (A) Schematic of dual-site recordings from HD cells in anterodorsal thalamic nuclei and retrosplenial cortex. (B) Behavioral protocol for in-vivo recordings involving sequences of two standard light sessions in the presence of a proximal cue (L_1_-L_2_), a 90° counterclockwise cue rotation session (R_3_), another standard light session (L_4_) followed by a dark session with cue removal (D_5_) and a final standard light session (L_6_). During rest, mice were moved on a flower pot. (C and G) Within-session stability of HD rate maps. HD cells from L7-PKCI mice exhibit significantly lower stability within the dark in both ADN (C, group × session, F_2.5, 386.6_ = 11.9, p < 0.001, two-way ANOVA, p < 0.001 post hoc Bonferroni) and RSC (G, group × session, F_2.7, 244.9_ = 5.1, p = 0.003, two-way ANOVA, p = 0.004 post hoc Bonferroni). (D and H) Relative per-cell changes of rate map stability compared to that of sessions L_1_. HD cells from both control and L7-PKCI have minor and similar changes in the standard light and cue-rotation sessions. In contrast, HD cells from L7-PKCI mice show a significant decrease within the dark in both ADN (D, group × session, F_2.4, 371.8_ = 12.1, p < 0.001, two-way ANOVA, p < 0.001 post hoc Bonferroni) and RSC (H, group × session, F_2.7, 240.8_ = 4.9, p = 0.004, two-way ANOVA, p = 0.007 post hoc Bonferroni). (E and I) Amount of HD rate map shift following the cue rotation session. HD cells from L7-PKCI mice followed the cue rotations in session R_3_ indifferent from controls in both ADN (E, group × session, F_1.7, 269.8_ = 0.8, p > 0.4, two-way ANOVA) and RSC (I, group × session, F_1.7, 153.3_ = 0.2, p > 0.7, two-way ANOVA). (F and J) Relative between-session similarity of HD rate maps in reference to a baseline similarity measured from L_2_×L_1_. HD rate maps in L7-PKCI mice exhibit significantly lower similarity of HD-rate maps between the dark (D_5_) and previous standard light session (L_4_) in ADN (F, group × session, F_1.2, 195.9_ = 9.4, p = 0.001, two-way ANOVA, p = 0.004, post hoc Bonferroni) and to a lesser extent in RSC (J, group × session, F_1.2,105.4_ = 5.6, p = 0.016, two-way ANOVA, p = 0.055 post hoc Bonferroni). (K and L) Representative HD cells from ADN (K) and RSC (L) in control (one cell, left columns) and L7-PKCI mice (two cells, middle and right columns). Each column represents the same cell across different sessions. The HD rate map and the HD×Time rate map are shown on the left and right for each cell, respectively. The inset values in black in the HD rate map represent the maximum firing rate of each cell (top) and inter-session stability measure (bottom). The inset values in red in the cue rotation session (R_3_) represent the amount of rate map shift this session compared to the previous standard light session (L_2_). The HD×Time rate maps represent the HD rate maps chunked in two-minute-long time bins. The inset value on top of each HDxTime rate map represents the maximum firing rate that occurred in a bin. The white inset line in the HD×Time rate map represents the mean preferred firing direction (PFD) of the cell in each time point. Note the change in the HD×Time rate map and the variation of PFD by time for L7-PKCI groups in both ADN (K) and RSC (L) within the dark session (D_5_).

We first analyzed the basic firing properties of identified HD cells from ADN and RSC in both control and L7-PKCI mutant mice. Spike amplitude, mean firing rate, and bursting index were all indistinguishable between the two groups in both structures (**Figures S3A and S3B**). Assessment of basic directional firing specificity, such as vector length and information, revealed that HD cells from L7-PKCI mice exhibit more substantial directional tuning than controls in the standard light sessions in ADN (**Figure S3C**) but not in RSC (**Figure S3D**). However, this difference was no longer present in the dark, during which control and mutant HD cells displayed indistinguishable directional properties. Finally, the peak firing rate of HD rate maps was not different between groups, suggesting that the HD cells from ADN of L7-PKCI mice did not fire more strongly but rather more exclusively at their PFD under light sessions (**Figure S3C and S3D**).

HD cells are known to sustain a highly stable preferred firing direction (PFD) in the presence of an external sensory cue, and to exhibit a drift by time from their PFD when the external sensory drive is removed (Goodridge et al., 1998). In the standard light and cue-rotation sessions, the stability of HD cells was comparable between L7-PKCI and control mice across all sessions in both ADN (**Figures 1C, 1K, and Figure S4;** group × session, F_2.5, 386.6_ = 11.9, p < 0.001, two-way ANOVA, p > 0.2 post hoc Bonferroni for L_1_, L_2_, L_4_, and L_6_) and RSC (**Figures 1G, 1L, and Figure S4;** group × session, F_2.7, 244.9_ = 5.1, p = 0.003, two-way ANOVA, p > 0.3 post hoc Bonferroni for L_1_, L_2_, L_4_, and L_6_). In the dark session, HD cells from both control and L7-PKCI mice exhibited a significant decrease in stability compared to the first light session in both ADN (**Figures 1C, 1K, and Figure S4;** group × session, F_2.5, 386.6_ = 11.9, p < 0.001, two-way ANOVA, p =0.026 post hoc Bonferroni for D_5_ versus L_1_ in control and p < 0.001 in L7-PKCI) and RSC (**Figures 1G, 1L, and Figure S4;** group × session, F_2.7, 244.9_ = 5.1, p = 0.003, two-way ANOVA, p =0.018 post hoc Bonferroni for D_5_ versus L_1_ in control and p < 0.001 in L7-PKCI). Interestingly however, HD cells from L7-PKCI mice showed significantly lower stability within the dark session compared to control in both ADN (**Figures 1C, 1K, and Figure S4;** group × session, F_2.5, 386.6_ = 11.9, p < 0.001, two-way ANOVA, p < 0.001 post hoc Bonferroni) and RSC (**Figures 1G, 1L, and Figure S4;** group × session, F_2.7, 244.9_ = 5.1, p = 0.003, two-way ANOVA, p = 0.004 post hoc Bonferroni). We also analyzed the relative changes of stability for each cell from the first light session and found out that HD cells from L7-PKCI and control mice have similar variation across light sessions in both ADN (**Figures 1D, and 1K;** group × session, F_2.4, 371.8_ = 12.1, p < 0.001, two-way ANOVA, p > 0.05 post hoc Bonferroni for L_2_, L_4_, and L_6_) and RSC (**Figures 1H, and 1L;** group × session, F_2.7, 240.8_ = 4.9, p = 0.004, two-way ANOVA, p > 0.3 post hoc Bonferroni for all L_2_, L_4_, and L_6_). However, the stability of HD cells from L7-PKCI mice dropped significantly in dark relative to the first light session in comparison to control mice in both structures (**Figures 1D, 1H, 1K and 1L;** ADN: group × session, F_2.4, 371.8_ = 12.1, p < 0.001, two-way ANOVA, p < 0.001 post hoc Bonferroni; RSC: group × session, F_2.7, 240.8_ = 4.9, p = 0.004, two-way ANOVA, p = 0.007 post hoc Bonferroni). We also assessed whether this effect was specific to any HD subclass but did not find any correlation between HD cell firing stability in the dark and cells’ basic firing characteristics or basic HD firing profile (**Figure S5**). Altogether, these results suggest that while control HD cells may be driven by the same gradient of multiple sensory drives across light and dark sessions, in L7-PKCI mice, a drastic change in processing the sensory drive in HD cells may occur when mice are conditioned to the dark session (**Figure S6).**

Next, we investigated the use of allocentric cues by the HD cells. In an environment with a salient external cue, it is known that the cue will be used as the reference frame by HD cells, potentially anchoring the internal representation of direction to the external world (Clark et al., 2012; Goodridge et al., 1998; Park et al., 2019; Taube et al., 1990b). Therefore, first, we tested whether HD rate maps follow a 90° counterclockwise rotation of a singular external cue within the arena (session R_3_, **Figure 1B**). Comparing HD rate maps from the 90° cue rotation session (R_3_) with the first light session (L_1_), we found a significant rotation of HD rate maps in both L7-PKCI and control mice (**Figures 1E, 1I, 1K, and 1L;** ADN: 85.5 ± 2.1° control, 79.9 ± 2.5° L7-PKCI, session main effect, F_1.7, 269.8_ = 1514.3, p < 0.001, two-way ANOVA; RSC: 83.7 ± 2.2° control, 79.8 ± 3.4° L7-PKCI, session main effect, F_1.7, 153.3_ = 1305.8, p < 0.001, two-way ANOVA) that fits the degree and direction of cue rotation. Meanwhile, the degree of rate map rotation was indistinguishable between control and L7-PKCI mice in both structures (ADN: group × session, F_1.7, 269.8_ = 0.8, p > 0.4; RSC: group × session, F_1.7, 153.3_ = 0.2, p > 0.7, two-way ANOVA). The rotation in HD rate maps was specifically observed during cue rotation sessions (R_3_) but not across standard light sessions, as illustrated by the shifts in the HD rate map clustered around zero in sessions L_2_ and L_4_ referenced to L_1_ in both controls (**Figures 1E, 1I, 1K, and 1L;** ADN: p > 0.9, post hoc Bonferroni, 3.1 ± 1.9° L_2_×L_1_, 2.9 ± 2.8° L_4_×L_1_; RSC: p > 0.9, post hoc Bonferroni, 1.3 ± 1.5° L_2_×L_1_, 0.1 ± 1.9° L_4_×L_1_) and L7-PKCI (**Figures 1E, 1I, 1K, and 1L;** ADN: p > 0.5, post hoc Bonferroni, 1.9 ± 1.2° L_2_×L_1_, - 0.3 ± 1.3° L_4_×L_1_; RSC: p > 0.9, post hoc Bonferroni, -0.9 ± 1.7° L_2_×L_1_, -1.7 ± 2.0° L_4_×L_1_). These data, therefore, indicate that HD cells from L7-PKCI mice are anchored to the allocentric cue properly and specifically impaired when relying on self-motion information.

To further investigate this idea, we analyzed more directly the similarity of HD rate maps across sessions. To this end, we compared the per-cell similarity of HD rate maps across light and dark sessions by using the similarity of HD rate maps from L_1_ and L_2_ as a baseline index for each cell (**Figures 1F and 1J**). The HD rate maps of control and L7-PKCI groups had indistinguishable similarity index (i.e. variation in the stability measure) across light sessions (**Figures 1F and 1J;** ADN: group × session, F_1.2, 195.9_ = 9.4, p = 0.001, two-way ANOVA, p > 0.05, post hoc Bonferroni; RSC: group × session, F_1.2, 105.4_ = 5.6, p = 0.016, two-way ANOVA, p > 0.05 post hoc Bonferroni). However, the HD rate map similarity index of L7-PKCI mice was significantly decreased from its baseline in the dark versus light session (**Figures 1F and 1J;** ADN: p < 0.001, post hoc Bonferroni for both D_5_×L_4_ versus L_6_×L_4_ and D_5_×L_4_ versus L_4_×L_2_; RSC: p < 0.001, post hoc Bonferroni for both D_5_×L_4_ versus L_6_×L_4_ and D_5_×L_4_ versus L_4_×L_2_) and also in comparison with the control mice (**Figures 1F and 1J;** ADN: p = 0.004, post hoc Bonferroni; RSC: p = 0.055, post hoc Bonferroni). Meanwhile, in the control group, the HD rate map similarity index of dark versus light session was statistically indistinguishable from its baseline (**Figures 1F and 1J;** ADN: p > 0.1, post hoc Bonferroni for both D_5_×L_4_ versus L_6_×L_4_ and D_5_×L_4_ versus L_4_×L_2_; RSC: p > 0.05, post hoc Bonferroni for both D_5_×L_4_ versus L_6_×L_4_ and D_5_×L_4_ versus L_4_×L_2_). Together, these results indicate that in L7-PKCI mice, but not in control mice, HD cells exhibit dissimilar rate maps in light and dark conditions. Such rate map modification from light to dark might arise from a drastic change in the type or in the weight of sensory inputs driving HD cells, potentially generating an altered representation in the dark or a new one that does not resemble the direction representation within the light. In support of this, while the stability score of control HD cells across light and dark sessions were correlated, such correlation was abolished in L7-PKCI mice in the dark (**Figure S6**).

### Impaired HD Cell Anchoring to Allocentric Cue in L7-PP2B Mice

We performed extracellular single-unit recordings from HD cells in ADN and RSC of six L7-PP2B mice and six control littermates in experimental conditions identical to L7-PKCI groups (**Figure 1A**, **Table S1, and Figures S7-S8).** The basic firing characteristics of HD cells (involving spike amplitude, mean firing rate, and burst index) and basic directional firing specificities (involving peak rate, mean vector length, directional information) were all similar between controls and L7-PP2B mice in both ADN and RSC structures (**Figure S9**).

Analyses of HD cells stability revealed that, under standard light conditions, HD cells from L7-PP2B mice were less stable than controls in ADN (**Figures 2A**; group × session, F_4.0, 717.5_ = 6.3, p < 0.001, two-way ANOVA, p < 0.05 for L_1_ and p < 0.01 for L_2_, L_4_, and L_6_, post hoc Bonferroni) but not in RSC (**Figures 2E**; group × session, F_2.7, 134.0_ = 1.9, p > 0.1, two-way ANOVA). This impaired stability was not observed in the dark (**Figures 2A and 2E**; ADN: group × session, F_4.0, 717.5_ = 6.3, p < 0.001, two-way ANOVA, p > 0.05, post hoc Bonferroni; RSC: group × session, F_2.7, 134.0_ = 1.9, p > 0.1, two-way ANOVA). Similar to HD cells from L7-PKCI mice and their respective control group, here in the RSC, HD cells from both L7-PP2B mice and their respective control group had a significant decrease in the stability within the dark session compared to the first light session (control: p < 0.001, post hoc Bonferroni; L7-PP2B: p = 0.009, post hoc Bonferroni). Interestingly, however, in the ADN, while control HD cells had a similar trend with a significant decrease in the stability within the dark session (p < 0.001, post hoc Bonferroni), HD cells from L7-PP2B exhibited indistinguishable stability between the dark and first light session (p > 0.9, post hoc Bonferroni). This suggests that, in ADN, individual HD cells from L7-PP2B mice but not from controls may retain the same stability in the dark as in the light (**Figure 2I**, the representative cell number 3 recorded from ADN of an L7-PP2B mouse exhibits a similar directional drift by time in both light and dark). To access this further, we analyzed the relative per-cell changes of stability across all sessions in reference to the first light session (**Figure 2B and 2F**). We found that HD cells from L7-PP2B and control mice have similar variation across light sessions in both ADN (**Figures 2B, and 2I;** group × session, F_2.1, 379.1_ = 6.2, p = 0.002, two-way ANOVA, p > 0.05, post hoc Bonferroni, for L_2_, L_4_, and L_6_) and RSC (**Figures 2F, and 2J;** group × session, F_2.4, 118.8_ = 2.0, p > 0.1, two-way ANOVA). However, in ADN, while the changes in the stability of HD rate maps in the control group were significantly lower in the dark compared to any other light session (**Figures 2B, and 2I**; p < 0.01, post hoc Bonferroni, for D_5_ versus L_2_, L_4_, and L_6_), no significant changes were observed between dark and any other light session for L7-PP2B mice (**Figures 2B, and 2I**; p > 0.05, post hoc Bonferroni, for D_5_ versus L_2_, L_4_, and L_6_). Further, in ADN, HD cells from L7-PP2B mice displayed significantly higher stability in the dark compared to the control group (**Figures 2B;** p < 0.001, post hoc Bonferroni).

**Figure 2.**
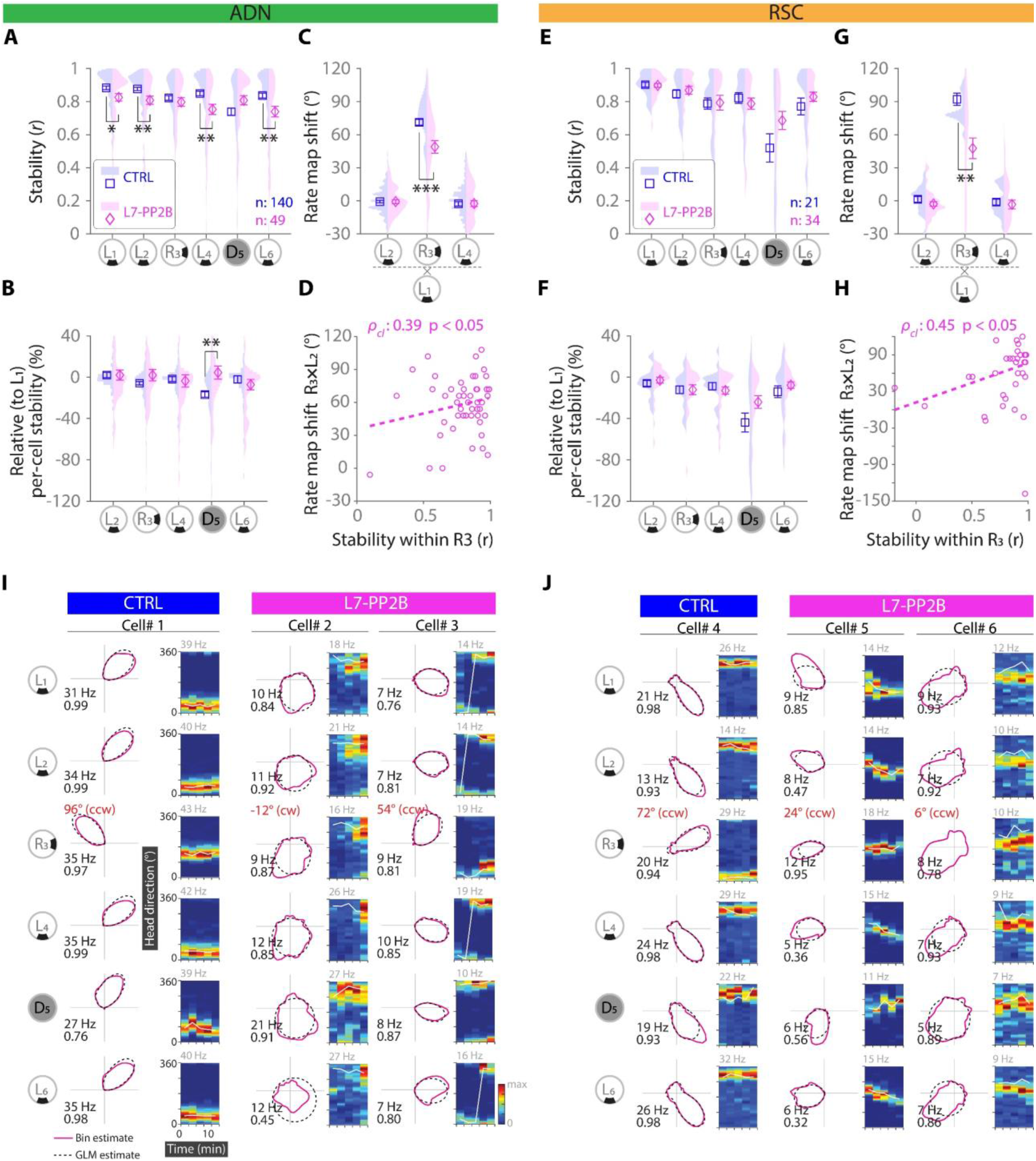
Impaired HD Cell Anchoring to Allocentric Cue in L7-PP2B Mice. (A and E) Within-session stability of HD rate maps. HD cells from L7-PP2B mice in ADN (A, group × session, F_4.0, 717.5_ = 6.3, p < 0.001, two-way ANOVA, p < 0.05 for L_1_ and p < 0.01 for L_2_, L_4_, and L_6_, post hoc Bonferroni) but not RSC (E, group × session, F_2.7, 134.0_ = 1.9, p > 0.1, two-way ANOVA) exhibit significantly lower stability within standard light sessions. (B and F) Relative per-cell changes of rate map stability compared to that of sessions L_1_. HD cells from both control and L7-PP2B have minor and similar changes in the standard light and cue-rotation sessions while HD cells in ADN (B, group × session, F_2.1, 379.1_ = 6.2, p = 0.002, two-way ANOVA, p < 0.001, post hoc Bonferroni) from L7-PP2B mice show significant higher values within the dark. (C and G) Amount of HD rate map shift following the cue rotation session. HD cells in L7-PP2B mice remained with a significantly lower average rotation in comparison to control in both ADN (C, group × session, F_1.7, 303.1_ = 15.6, p < 0.001, two-way ANOVA, p < 0.001, post hoc Bonferroni; 71.2 ± 2.6° control, 48.7 ± 5.9° L7-PP2B) and RSC (G, group × session, F_1.2, 63.0_ = 10.5, p = 0.001, two-way ANOVA, p = 0.001, post hoc Bonferroni; 92.1 ± 5.7° control, 47.6 ± 9.5° L7-PP2B). (D and H) The linear-directional correlation analysis of HD cells from L7-PP2B mice revealed that the amount of rate map shift following the cue rotation and the rate map stability within the cue rotation session (R_3_) are positively correlated in both ADN (D, p*_cl_* =0.39, p < 0.05 circular-linear Pearson correlation) and RSC (H, p*_cl_* =0.45, p < 0.05 circular-linear Pearson correlation). (I and J) Representative HD cells from ADN (I) and RSC (J) in control (one cell, left columns) and L7-PKCI mice (two cells, middle and right columns). Each column represents the same cell across different sessions. The HD rate map and the HD×Time rate map are shown on the left and right for each cell, respectively. The inset values in black in the HD rate map represent the maximum firing rate of each cell (top) and inter-session stability measure (bottom). The inset values in red in the cue rotation session (R_3_) represent the amount of rate map shift this session compared to the previous standard light session (L_2_). The HD×Time rate maps represent the HD rate maps chunked in two-minute-long time bins. The inset value on top of each HDxTime rate map represents the maximum firing rate that occurred in a bin. The white inset line in the HD×Time rate map represents the mean preferred firing direction (PFD) of the cell in each time point. In the L7-PP2B group, Note the lower stability of HD×Time rate maps within light sessions (L_1_, L_2_, L_4_, L_6_), the high correlation values comparable to that in the light session in ADN (I), and the weak HD rate map PFD shifts following cue rotation (R_3_) in both ADN (I) and RSC (J).

During cue-rotation sessions, both groups had a significant HD rate map shift compared to the other light sessions (**Figures 2C, 2G, 2I, and 2J;** ADN: group × session, F_1.7, 303.1_ = 15.6, p < 0.001, two-way ANOVA, p < 0.001 post hoc Bonferroni for both R_3_×L_1_ versus L_2_×L_1_ and R_3_×L_1_ versus L_4_×L_1_ comparisons; RSC: group × session, F_1.2, 63.0_ = 10.5, p = 0.001, two-way ANOVA, p < 0.001 post hoc Bonferroni for both R_3_×L_1_ versus L_2_×L_1_ and R_3_×L_1_ versus L_4_×L_1_ comparisons). However, while a nearly 90° counterclockwise rotation of rate maps following the cue rotation was observed, as expected, in control mice, HD rate maps from L7-PP2B mice rotated with a significantly lower average degree in both ADN (**Figures 2C, and 2I;** p < 0.001, post hoc Bonferroni; 71.2 ± 2.6° control, 48.7 ± 5.9° L7-PP2B) and RSC (**Figures 2G, and 2J;** p = 0.001, post hoc Bonferroni; 92.1 ± 5.7° control, 47.6 ± 9.5° L7-PP2B). Further, we found that in L7-PP2B mice, the amount of HD rate map shift was significantly correlated with HD rate map stability within the cue-rotation sessions in both ADN (**Figures 2D and 2I**; p*_cl_* =0.39, p < 0.05 circular-linear Pearson correlation) and RSC (**Figures 2H and 2J**; p*_cl_* =0.45, p < 0.05 circular-linear Pearson correlation), indicating that the cells that are less strongly driven by the external cue exhibit less stable HD rate maps within the cue rotation session. Finally, HD rate map similarity across different standard light sessions was indistinguishable in L7-PP2B mice and control, as shown by the shifts in the HD rate map in sessions L_2_ and L_4_ in reference to session L_1_ that was clustered around zero in both controls (**Figures 2C, 2G, 2I, and 2J;** ADN: p > 0.7, post hoc Bonferroni, 3.1 ± 1.9° L_2_×L_1_, 2.9 ± 2.8° L_4_×L_1_; RSC: p > 0.9, post hoc Bonferroni, 1.3 ± 1.5° L_2_×L_1_, 0.1 ± 1.9° L_4_×L_1_) and L7-PP2B mice (ADN: p > 0.9, post hoc Bonferroni, 1.9 ± 1.2° L_2_×L_1_, -0.3 ± 1.3° L_4_×L_1_; RSC: p > 0.9, post hoc Bonferroni, -0.9 ± 1.7° L_2_×L_1_, -1.7 ± 2.0° L_4_×L_1_). Together, these results indicate that under the light conditions, while the allocentric cue strongly drives HD cells from control mice, this allocentric drive was weaker in L7-PP2B mice. Further, in contrast to the expected decrease observed in control mice in the HD rate maps stability from light to dark sessions, potentially due to the error introduced by the path integration process, HD rate maps of L7-PP2B mice maintained a similar rate map stability from light to dark conditions. This data, together with the lower stability observed in standard light conditions compared to controls, suggests that the external versus idiothetic inputs ratio driving HD cell firing may have been affected in L7-PP2B mice, with an increase in the idiothetic drive over the external one.

### Independent HD Cell Attractor Networks Are Coordinated by Cerebellar Mechanisms

We then questioned whether the sub-second temporal correlation structure is preserved under different sensory conditions between HD cell pairs, both within and between ADN and RSC structures, and under cerebellar alterations.

Under standard light and cue-rotation sessions, pairs of simultaneously recorded HD cells from both L7-PKCI, L7-PP2B, and their respective control littermates showed a preserved correlation structure across different sessions (**Figure 3 and Figure S10;** p > 0.2 for all conditions, two-sided Z-test, Bonferroni correction for multiple comparisons). In the dark session, however, the correlation structure of HD cell pairs from L7-PKCI mice was significantly decreased compared to its baseline (**Figure 3 and Figure S10;** p < 0.001, two-sided Z-test, Bonferroni correction for multiple comparisons), while it was preserved for all other three groups of mice (control littermates of L7-PKCI as well as L7-PP2B mice and their control littermates; **Figure 3 and Figure S10;** p > 0.2 for all conditions, two-sided Z-test, Bonferroni correction for multiple comparisons). Expectedly, the correlation structure for these three groups was preserved both within ADN, within RSC, and also between ADN and RSC HD cell pairs, across all sessions (**Figures S11, S12, and S13;** p > 0.3 for all conditions, two-sided Z-test, Bonferroni correction for multiple comparisons).

**Figure 3.**
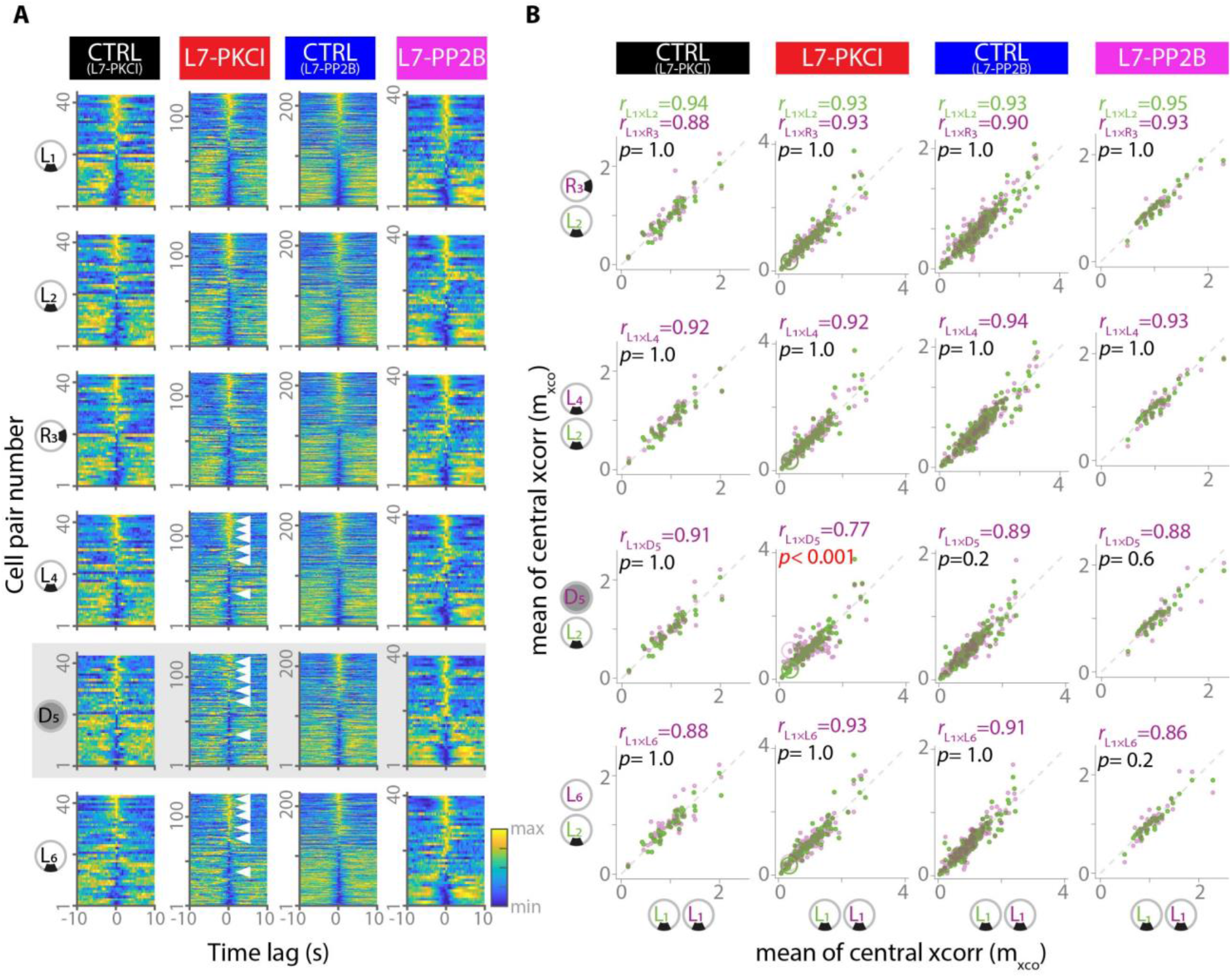
Impaired Temporal Coordination between HD Cells in L7-PKCI Mice when Relying on Self-Motion Cues and Preserved Temporal Coordination in L7-PP2B Mice. (A) Temporal cross-correlograms of all simultaneously recorded HD cell pairs for all four groups of mice across the six sessions. Within each heat-map, each row represents the cross-correlogram of one HD cell (B) pair. The HD cell pairs are ordered in all sessions based on their m_xco_ value (mean of cross-correlogram in central two-second bins) from session L_1_, with the highest m_xco_ value in session L_1_ in the topmost row to the lowest m_xco_ value in the bottom-most row. In the L7-PKCI group, white triangular markers are for visual aid in comparing exemplar pairs between the dark session (D_5_) and the standard light sessions before (L_4_) and after (L_6_) the dark. (C) Pearson correlation between m_xco_ value of each session and session L_1_. In each comparison, the correlation between session L_2_ and L_1_ (green) is used as the baseline, and the correlation between m_xco_ value of any other session and session L_1_ (purple) is compared to this baseline. Only HD cell pairs in L7-PKCI mice and only within the dark sessions exhibited a correlation significantly lower than their baseline from all the groups and all conditions. For the L7-PKCI group, one representative cell pair is highlighted by a larger circle for visual comparisons between different conditions. The inset value in each scatter plot represents the correlation value between L_1_ and L_2_ (top, green), the correlation value between L_1_ and the session indicated in purple (middle, purple), and the p-value from comparing the two correlations (green versus purple distribution). The *p*-Values are after the two-sided Z-test, with Bonferroni correction for multiple comparisons. The m_xco_ values of one exemplar cell pair are highlighted by the large green and purple circles for the L7-PKCI group. The two circles deviate from each other, specifically in the dark.

Interestingly, however, the impaired correlation structure in L7-PKCI mice within the dark session was specific to the HD cell pairs between ADN and RSC (**Figure 4**; p < 0.001 for ADN×RSC in dark session, two-sided Z-test, Bonferroni correction for multiple comparisons). Indeed, HD cell pairs within ADN and within RSC showed a preserved correlation structure across all sessions (**Figure 4F and 4G**; p > 0.9 for all sessions, two-sided Z-test, Bonferroni correction for multiple comparisons). Further, the correlation structure in L7-PKCI mice between ADN and RSC HD cell pairs was preserved across standard light and cue-rotation sessions (**Figure 4F and 4G**; p > 0.9 for all sessions, two-sided Z-test, Bonferroni correction for multiple comparisons), suggesting that the coordination of activity between HD cell pairs from RSC and ADN is altered in L7-PKCI mice specifically when a salient external sensory drive is lacking.

**Figure 4.**
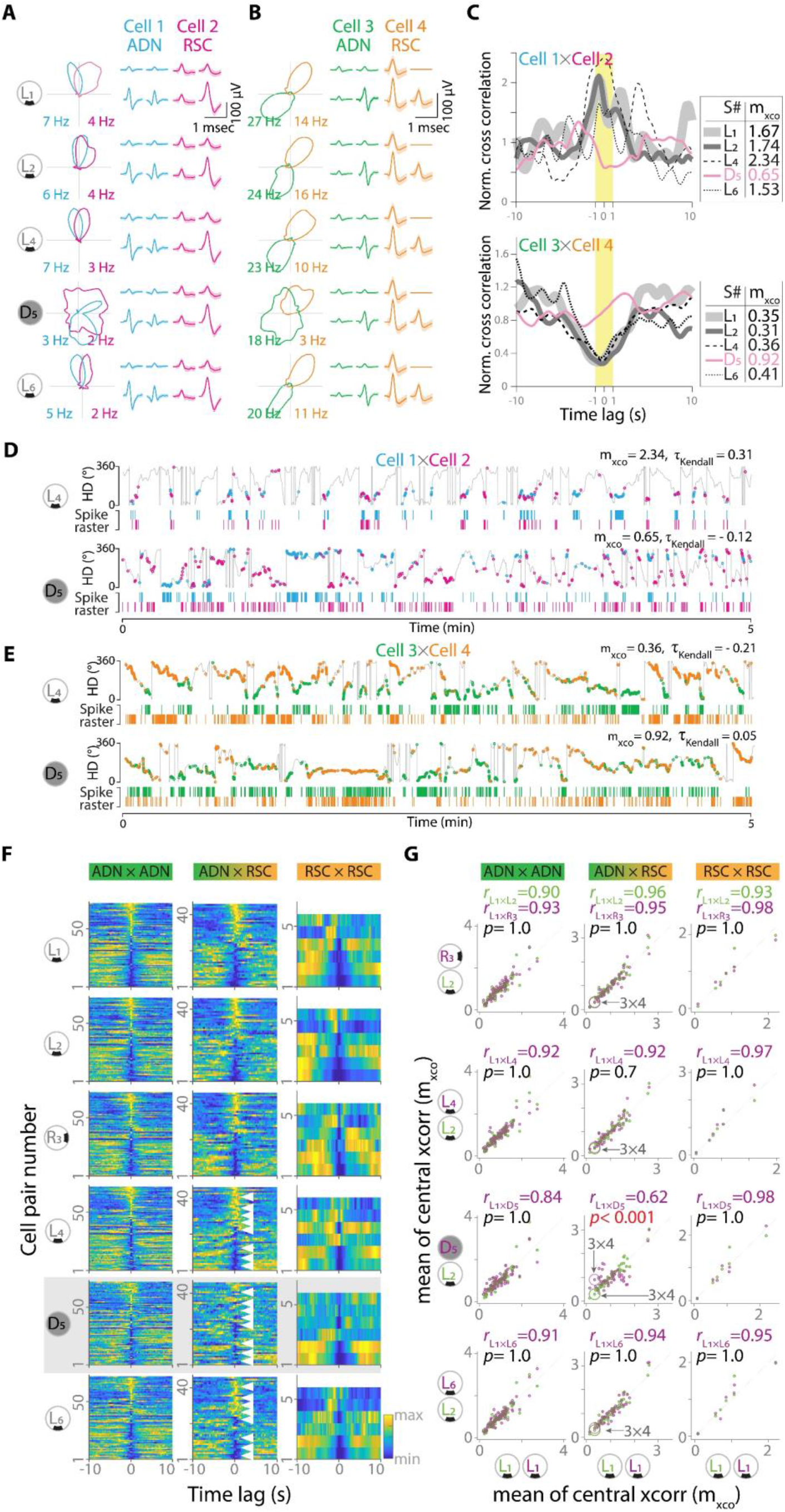
Impaired Temporal Coordination of L7-PKCI HD Cells is Specific between ADN and RSC Networks. (A-B) The HD rate maps (left) and spike waveforms (middle and right) for representative HD cell pairs, pair 1×2 (A) and pair 3×4 (B) recorded from L7-PKCI mice. Cells 1 and 3 were recorded from ADN, and cells 12 and 4 were recorded from RSC. Spike waveforms are represented as mean±sem. (A) Temporal cross-correlogram of HD cell pairs 1×2 (top) and HD cell pair 3×4 (bottom) within the standard light and dark sessions. Note the cross-correlation changes in the central 2-second (Yellow rectangular box) within the dark session for both cell pairs while remaining similar across different light sessions. m_xco_ represents the mean value of the central 2-second temporal cross-correlogram. (D-E) A 5-min sample of spike raster along the animal head direction for cell pair 1×2 (E) and 3×4 (F) within the standard light session L_4_ (top) and the dark session D_5_ (bottom). With an overlapping HD rate map, the HD cell pair 1×2 exhibits a high temporal correlation through spike co-discharge in a short-timescale within the standard light sessions (positive Kendall’s correlation). However, they fire in separate time windows within the dark session (negative Kendall’s correlation). With a separated PFD, the HD cell pair 3×4 exhibits a low temporal correlation by firing in separate time windows within the standard light session. However, they often co-fire within the dark session. τ_Kendall_ represents Kendall’s correlation between the spike timestamps of the pair measured at 1000-ms timescale (Supplemental Experimental Procedures) used as an alternative non-parametric approach for measuring the temporal correlation of cell pair. (F) Temporal cross-correlograms of all simultaneously recorded HD cell pairs in L7-PKCI mice within ADN (left), between ADN and RSC (middle), and within RSC (right) across all sessions. Within each heat-map, each row represents the cross-correlogram of one HD cell pair. For the ADN×RSC group, white triangular markers are for visual aid in comparing exemplar pairs between the dark session (D_5_) and the standard light sessions before (L_4_) and after (L_6_) the dark.. (G) Pearson correlation between m_xco_ value of each session and session L_1_ indicates that the correlation structure between ADN and RSC cells (middle) is specifically affected within the dark session. The correlation structure within each of ADN (top) and RSC (bottom) has remained intact. In each comparison, the correlation between session L_2_ and L_1_ (green) is used as the baseline. The correlation between m_xco_ value of any other session and session L_1_ (purple) is compared to this baseline. The inset value in each scatter plot represents the correlation value between L_1_ and L_2_ (top, green), the correlation value between L_1_ and the session indicated in purple (middle), and the p-value from comparing the two correlations (green versus purple). The *p*-Values are after the two-sided Z-test, with Bonferroni correction for multiple comparisons.

### Cerebellum Modulates HD Cell Firing Independently from Speed and AHV Codes

Along with direction information, thalamocortical HD cell systems are known to contain cells with activity patterns related to other navigational parameters such as linear speed and angular head velocity. Akin to HD cells, AHV, and speed coding cells are also known to be driven by a combination of vestibular and visual input components. Therefore, we tested whether cerebellar manipulations impact speed and AHV coding in those structures. The relative changes of speed score across the different light sessions and from light to dark session were similar between the two groups of cerebellar mutant mice and their respective control littermates, in both ADN (**Figures 5A, 5B, 6A and 6B**; L7-PKCI: group × session, F_1.7, 158.2_ = 1.35, p = 0.26, two-way ANOVA; L7-PP2B: group × session, F_2.8, 299.6_ = 0.49, p = 0.68, two-way ANOVA) and RSC (**Figures 5A and 6A**; L7-PKCI: group × session, F_1.5, 42.2_ = 2.93, p = 0.078, two-way ANOVA; L7-PP2B: group × session, F_2.5, 35.7_ = 0.88, p = 0.45, two-way ANOVA).

**Figure 5.**
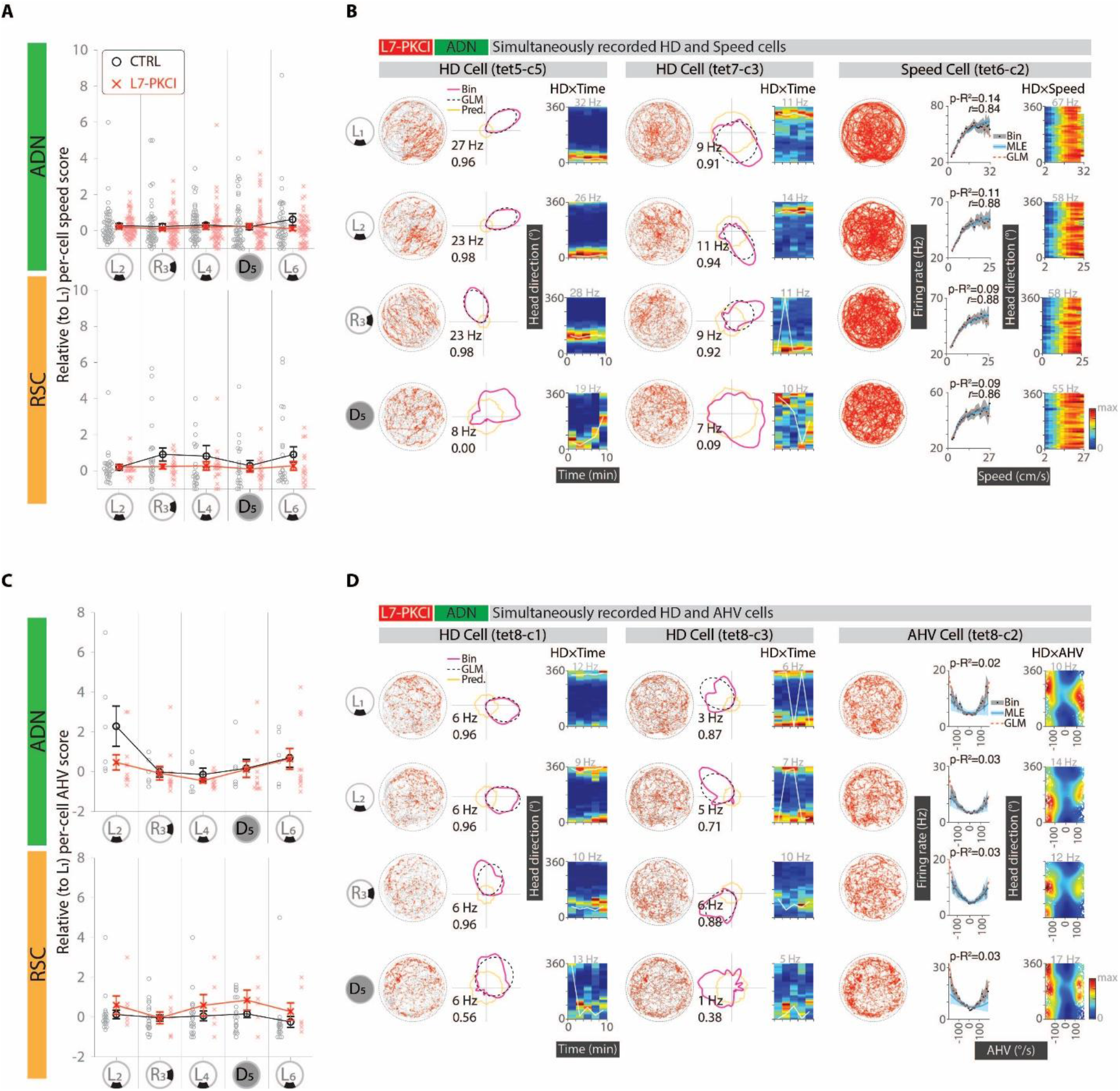
Intact Linear Speed and Angular Head Velocity Coding in Thalamocortical HD Cell System in L7-PKCI mice. (A) Changes of speed score for sessions L_2_ to L_6_ relative to the first standard light session, L_1_. Speed score changes were indifferent between L7-PKCI and control in both ADN (left, group × session, F_1.7, 158.2_ = 1.35, p = 0.26, two-way ANOVA) and RSC (right, group × session, F_1.5, 42.2_ = 2.93, p = 0.078, two-way ANOVA). (B) Representative simultaneously recorded HD (left and middle) and speed cells (right) from the ADN structure of L7-PKCI mice. While the firing of both HD cells becomes unstable within the dark session, speed cell firings remain stable, similar to that in light sessions. For each HD cell, trajectory map (left; animal tracking on gray and spike in red dots; Inset values represent maximum firing rate on top and stability measure in the bottom), HD rate maps (middle; bin estimate in magenta, GLM estimate in black and predicted estimate in orange) and HD×Time rate map (right, Inset gray value on top represent maximum firing) are shown. For speed cell, trajectory map (left), speed rate map (middle; bin estimate in gray, MLE estimate in cyan, and GLM estimate in red; Inset values represent pseudo-R^2^ on top and Pearson’s correlation coefficient *r*), and HD×Speed rate map (right, Inset gray value on top represent maximum firing) are shown. (C) Changes of AHV score for sessions L_2_ to L_6_ relative to the first standard light session, L_1_. AHV score changes were indifferent between L7-PKCI and control in both ADN (left, group × session, F_2.7, 41.2_ = 2.05, p = 0.13, two-way ANOVA) and RSC (right, group × session, F_4, 96_ = 0.43, p = 0.79, two-way ANOVA). (D) Representative simultaneously recorded HD (left and middle) and AHV cells (right) from the ADN structure of L7-PKCI mice. While the firing of both HD cells becomes unstable within the dark session, AHV cell firings remain stable similar to that in the light session. For AHV cell, trajectory map (left), AHV rate map (middle; bin estimate in gray, MLE estimate in cyan, and GLM estimate in red; Inset values indicated as p-R^2^ refers to pseudo-R^2^), and HD×AHV rate map (right, Inset gray value on top represent maximum firing) are shown.

**Figure 6.**
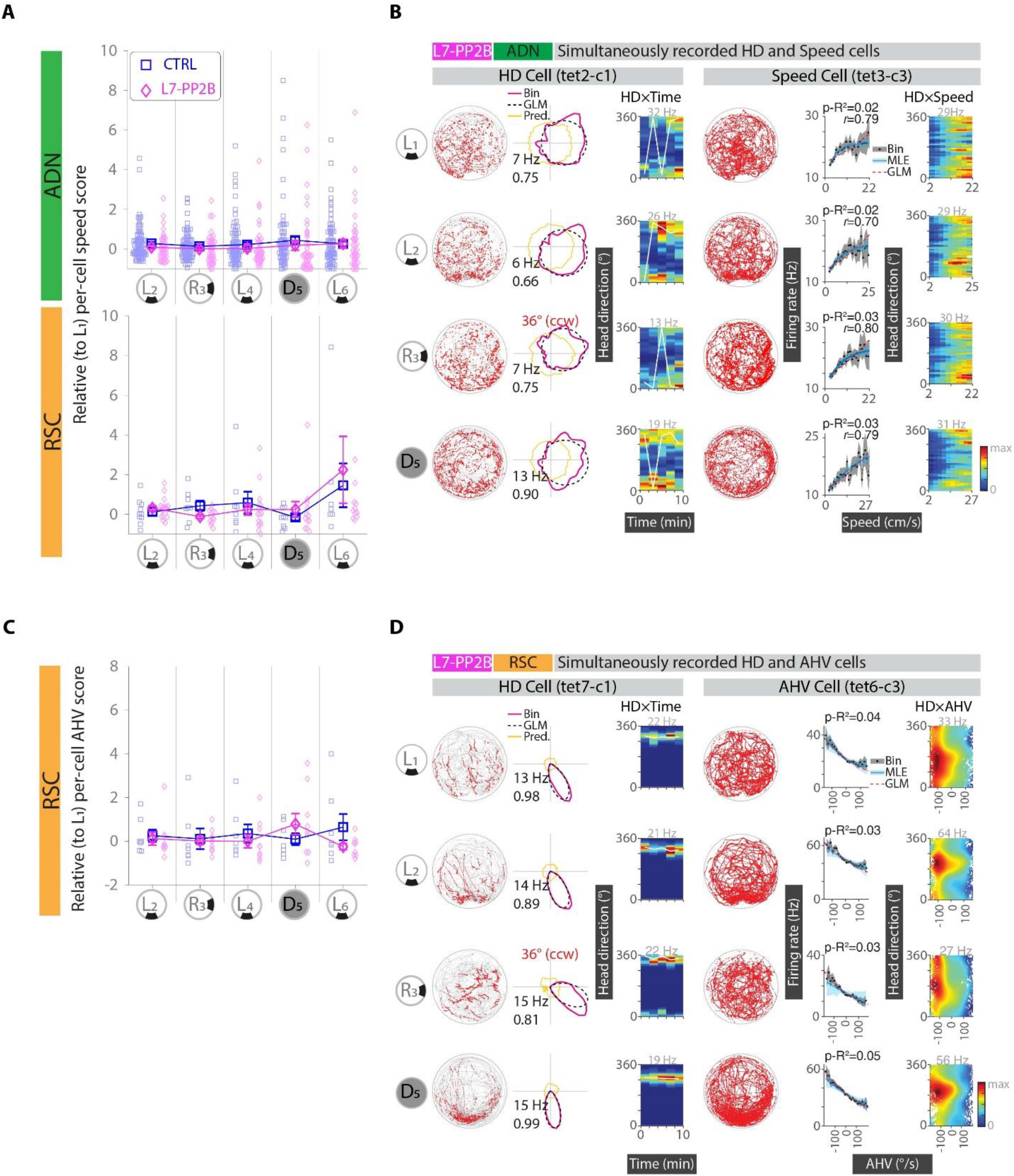
Intact Linear Speed and Angular Head Velocity Coding in Thalamocortical HD Cell System in L7-PP2B mice. (A) Changes of speed score for sessions L_2_ to L_6_ relative to the first standard light session, L_1_. Speed score changes were indifferent between L7-PP2B and control in both ADN (left, group × session, F_2.8, 299.6_ = 0.49, p = 0.68, two-way ANOVA) and RSC (right, group × session, F_2.5, 35.7_ = 0.88, p = 0.45, two-way ANOVA). (B) Representative simultaneously recorded HD (left) and speed cells (right) from the ADN structure of L7-PP2B mice. While HD cell firings are drifting within the light session, speed cell firings remain stable across all sessions. For each HD cell, trajectory map (left; animal tracking on gray and spike in red dots; Inset values represent max firing rate on top and stability measure in the bottom), HD rate maps (middle; bin estimate in magenta, GLM estimate in black and predicted estimate in orange) and HD×Time rate map (right, Inset gray value on top represent maximum firing) are shown. For speed cell, trajectory map (left), speed rate map (middle; bin estimate in gray, MLE estimate in cyan, and GLM estimate in red; Inset values represent pseudo-R^2^ on top and Pearson’s correlation coefficient *r*), and HD×Speed rate map (right, Inset gray value on top represent maximum firing) are shown. (C) Changes of AHV score for sessions L_2_ to L_6_ relative to the first standard light session, L_1_. AHV score changes were indifferent between L7-PP2B and control in RSC (group × session, F_2.5, 35.5_ = 1.51, p = 0.23, two-way ANOVA). Comparisons in ADN were not possible due to the insufficient number of recorded AHV cells. (D) Representative simultaneously recorded HD (left) and AHV cells (right) from the RSC structure of L7-PP2B mice. While the stability of HD cells varies between sessions and HD cell firing follows the cue rotation weakly, AHV cell firing remains stable across sessions. For AHV cell, trajectory map (left), AHV rate map (middle; bin estimate in gray, MLE estimate in cyan, and GLM estimate in red; Inset values indicated as p-R^2^ refers to pseudo-R^2^), and HD×AHV rate map (right, Inset gray value on top represent maximum firing) are shown.

Regarding AHV cells, here again, no difference was found between cerebellar mutant mice and their respective control littermate in the relative changes of AHV score across the different light sessions and from light to dark sessions in both ADN (**Figures 5C, 5D**; L7-PKCI: group × session, F_2.7, 41.2_ = 2.05, p = 0.13, two-way ANOVA; L7-PP2B: not enough cells) and RSC (**Figures 5C, 6C, and 6D**; L7-PKCI: group × session, F_4,96_ = 0.43, p = 0.79, two-way ANOVA; L7-PP2B: group × session, F_2.5, 35.5_ = 1.51, p = 0.23, two-way ANOVA).

Importantly, in the recordings during which AHV or speed cells were concomitantly recorded with HD cells, we observed a preserved speed and AHV coding when HD firing was altered in the dark for the L7-PKCI mice (**Figures 5B and 5D**, ADN), or in the light for the L7-PP2B mice, (**Figures 6B**, ADN; **Figure 6D**, RSC). These examples are particularly important as HD cells in ADN are suggested to be part of an intrinsic attractor network in which a change in one HD cell firing resembles the local network behavior. Altogether, these data suggest that altered HD firing in cerebellar mutant mice is not caused by an impaired AHV or speed coding in the recorded structures.

## DISCUSSION

By simultaneous recordings from HD cells of ADN and RSC structures of freely-behaving cerebellar-specific transgenic mouse models under different sensory conditions, we demonstrated: 1) the existence of sub-second temporal coordination between thalamic and retrosplenial HD cells that contribute to forming a unitary representation of direction 2) the role of cerebellar PKC-dependent mechanisms in maintaining this temporal coordination when the external sensory drive is removed and 3) the implication of distinct cerebellar mechanisms required for stable direction representation. These results put forward a new role for the cerebellum for maintaining a stable and unitary thalamocortical representation of direction under different sensory conditions.

Our finding in control mice shows that temporal coordination within ADN, RSC, and between ADN and RSC HD cell pairs is preserved in both light and dark conditions, suggesting a common HD attractor network within and between these brain structures.

Previous studies have suggested that both sensory signals and internal mechanisms are involved in coordinating neuronal activity in the HD system (Peyrache et al., 2015; Taube, 2007; Yoganarasimha et al., 2006; Zugaro et al., 2003). However, no input extrinsic to the HD system had ever been identified in mediating such network behavior. Therefore, it was unknown whether the unitary representation of direction is intrinsic to and self-organized by the HD system (Redish et al., 1996; Skaggs et al., 1995; Zhang, 1996). We show that the correlation structure between ADN and RSC cell pairs in L7-PKCI mice is specifically abolished upon removing external sensory signals. These results suggest that neuronal coordination between the ADN and RSC HD cells requires a stable prominent external sensory signal or cerebellar PKC-dependent processing. This suggests a role for the cerebellum in providing necessary internal computations in the absence of external sensory signals. To our knowledge, this is the first direct experimental demonstration of the existence of multiple independent HD attractor networks. The coordination of such separate HD networks is potentially fundamental for building a single and unique direction representation across thalamocortical brain structures (Hulse and Jayaraman, 2020). Our findings are the first to identify a role for the cerebellum in mediating neuronal coordination between distant HD structures.

Despite the altered neuronal coordination between ADN and RSC HD cell pairs of L7-PKCI mice under the dark, such coordination within each of these structures remained preserved. This within-structure neuronal coordination suggests a local, intrinsic, and potentially self-organized HD network within both ADN and RSC (or a more extensive network each in part belongs to) independent of both external sensory drive and internal cerebellar PKC- and PP2B-dependent mechanisms.

In addition to its role in coordinating HD cell activity among different regions, we found that cerebellar processes are also involved in stable direction representation. We found that HD cells from L7-PKCI mice exhibit rapidly drifting directional firing patterns under dark conditions. At the same time, their activity was similar to controls under light and cue rotation conditions. These results reinforce previous works highlighting the critical role of cerebellar PKC-dependent mechanisms in self-motion information processing during spatial navigation (Hernández et al., 2020; Rochefort et al., 2011). Unlike the L7-PKCI mice, HD cells from L7-PP2B mice exhibited a different alteration profile, with lower stability of directional firing pattern under light conditions and a lower control by prominent allocentric cue. Further, unlike controls, L7-PP2B HD cells sustained a more similar degree of directional stability from light to dark sessions. This result indicates that the potential input driving HD cell activity in L7-PP2B mice under the light has not been significantly changed when mice underwent dark conditions (with cue removal). Together these data suggest that in mice lacking cerebellar PP2B-dependent processes, idiothetic cues may predominantly influence HD rate maps in both dark and light conditions. One important note is that the idiothetic cues may not be the sole drive of L7-PP2B HD cells firing since the HD cells still follow cue rotations, albeit significantly weaker than control. These results from L7-PP2B, along with the results from L7-PKCI group, support the hypothesis that HD cells from wild-type mice are simultaneously driven by the integration of multimodal cues, even when a prominent visual cue is available (Savelli and Knierim, 2019). In contrast, in both cerebellar mutant mice, such multi-sensory integration is impaired, leading to less stable HD signals. These results fit with our hypothesis that the cerebellum is fueling the navigation circuitry with an appropriate sensory drive as the cerebellum is adequately wired to combine the diversity of sensory signals (Rondi-Reig et al., 2014) and PKC/PP2B-dependent mechanisms appear as molecular mechanisms at the core of these cerebellar computations toward coordinating neuronal activity between distant structures of the navigation circuit during behavior.

Interestingly, the contrasting results between L7-PKCI and L7-PP2B mice reported here are supporting our previous findings on hippocampal place cells that in L7-PP2B were partially unstable under the light (yet preserved in the dark), and in L7-PKCI were specifically unstable under dark conditions (Lefort et al., 2019; Rochefort et al., 2011). Previous studies performing lesions on head direction circuitry have reported severe spatial behavior deficits (Aggleton and Nelson, 2015), suggesting that HD system plays a role in hippocampal functions. Such lesions do not abolish but lead to unstable hippocampal place cell activity (Calton et al., 2003). Since such lesions have been performed on the network encompassing -but not specific to- HD cells, it was not known whether unstable hippocampal place coding was explicitly due to impaired HD cells function *per se*. The convergence of the effects presented here (specific alteration to HD cell coding but not AHV and speed coding) associated with our previous data on hippocampal place cells (Lefort et al., 2019; Rochefort et al., 2011) suggest that this might indeed be the case.

These results fit with our hypothesis that the cerebellum is fueling the navigation circuitry with an appropriate sensory drive as the cerebellum is adequately wired to combine the diversity of sensory signals (Rondi-Reig et al., 2014) and PKC/PP2B-dependent mechanisms appear as molecular mechanisms at the core of these cerebellar computations toward coordinating neuronal activity between distant structures of the navigation circuit during behavior.

Since the cerebellum is reciprocally connected with the vestibular system, one might interrogate if the altered thalamocortical temporal coordination in L7-PKCI mice and altered HD cell stability in both mutant mice can be due to the presence of a biased or decoupled angular velocity signal, leading to error accumulation in the path integration process (Goodridge et al., 1998; Green et al., 2017; Leonhard et al., 1996; Mittelstaedt and Mittelstaedt, 1980; Turner-Evans et al., 2017). Our observation that firing properties of AHV cells remained unchanged while the stability of HD cells in L7-PKCI mice under dark and L7-PP2B mice under light was lower (**Figure 5D**) excludes this idea. Instead, our data suggest that altered HD cell firing properties are caused by a deficit in cerebellar-dependent integration of multi-sensory signals. Further in this respect, a recent study characterizing the potential vestibular-driven (otolith and semicircular canals) Purkinje cell responses in the posterior vermis of the cerebellar cortex in L7-PKCI mice suggests that cerebellar PKC-dependent mechanisms modulate the gain of translation information at the single Purkinje cell level (Hernández et al., 2020). Whether this effect is specific to the linear or angular acceleration component of translation was not addressed, albeit the directional tuning of neuronal responses were similar between groups. In line with their finding, we found a more substantial linear speed coding within HD circuitry in both ADN and RSC structures of L7-PKCI mice (**Figure S15**). Importantly, however, the speed coding did not change under the dark condition in which the alteration in direction coding of concomitant HD cells was observed. Therefore, it is unlikely that specific alteration of HD rate-maps in the dark is related to a difference in computation of linear speed signal presented at the speed cell level and potentially the acceleration components in the translation information encoded by Purkinje cells in the posterior vermis. Another support on the reasoning that in L7-PKCI mice, idiothetic cues are not optimally integrated into the spatial system comes from firing specificity of HD and place cells (Rochefort et al., 2011) recorded from this mouse model under standard light conditions. Thalamic HD cells and hippocampal place cells exhibited significantly stronger directional (**Figure S9**) and spatial specificity (Rochefort et al., 2011), respectively. In standard light conditions in which external cues are present, the idiothetic cues are continuously monitored and integrated with existing external sensory information (Jayakumar et al., 2019; Savelli and Knierim, 2019). Our results suggest that such integration is lacking in L7-PKCI mice resulting in HD and place cells to be driven almost exclusively by external sensory information. In line with this view, while control HD cells showed correlated stability scores across light and dark sessions, such correlation was abolished in L7-PKCI HD cells under the dark, suggesting a more drastic change in the driving input under dark conditions compared to control (**Figure S6**).

Finally, along with HD cells, we found a prominent speed signal in the thalamocortical HD system. We observed very few theta-rhythmic HD and speed cells in ADN structure and almost no theta-rhythmic HD or speed cell in RSC of all four mice groups (**Figures S3, S9 and S15**). These results support the idea that the thalamocortical HD network and their associated speed signal may function parallel to the septal oscillatory inputs arriving limbic system (Dillingham and Vann, 2019). This hypothesis is in line with studies dissociating the speed and oscillatory signal within the medial entorhinal cortex, a structure downstream to ADN and RSC (Hinman et al., 2016). However, the speed and oscillatory signals are intermingled within neuronal firings of the medial entorhinal cortex (Hinman et al., 2016; Kropff et al., 2015). Therefore, our results suggest that similar to the HD signal (Winter et al., 2015), the non-oscillatory speed signal in the HD circuitry reported here might be afferent to that of the medial entorhinal cortex. The specific effects of cerebellar alteration to HD but not speed coding raises whether the speed signal in the HD circuitry has a separate neural substrate and functions in parallel to the HD cells, which needs to be examined in the future.

In summary, our data suggest the existence of multiple HD attractor networks in the mammalian brain in which their coordination toward building a single and stable direction representation is facilitated by different cerebellar mechanisms.

## EXPERIMENTAL PROCEDURES

Experimental procedures are covered in Supplemental Experimental Procedures.

## SUPPLEMENTAL INFORMATION

Supplemental Information includes Supplemental Experimental Procedures, fifteen figures, and one table which can be found with this article online.

## AUTHOR CONTRIBUTIONS

Conceptualization: MF and LRR; Methodology: MF, CR, and LRR; Experiment Setup: MF; Experimentation: MF, JLM, XZ, JV; Software, Investigation, and Analysis: MF; Validation: MF, CR, and LRR; Visualization, MF; Writing Original Draft, Review, and Editing: MF, CR, and LRR; Supervision, Funding Acquisition, and Project administration: LRR.

## ACKNOWLEDGMENTS

This work was supported by the Fondation pour la Recherche Médicale DEQ20160334907-France and by the National Agency for Research ANR-17-CE37-0015-01-NaviGPS (LRR), CNRS, Inserm and Sorbonne Université. We thank all members of the CEZAME team for helpful discussions of the experiments and manuscript. We gratefully acknowledge the IBPS animal facility staff for their support.

---

**Figure S1.**
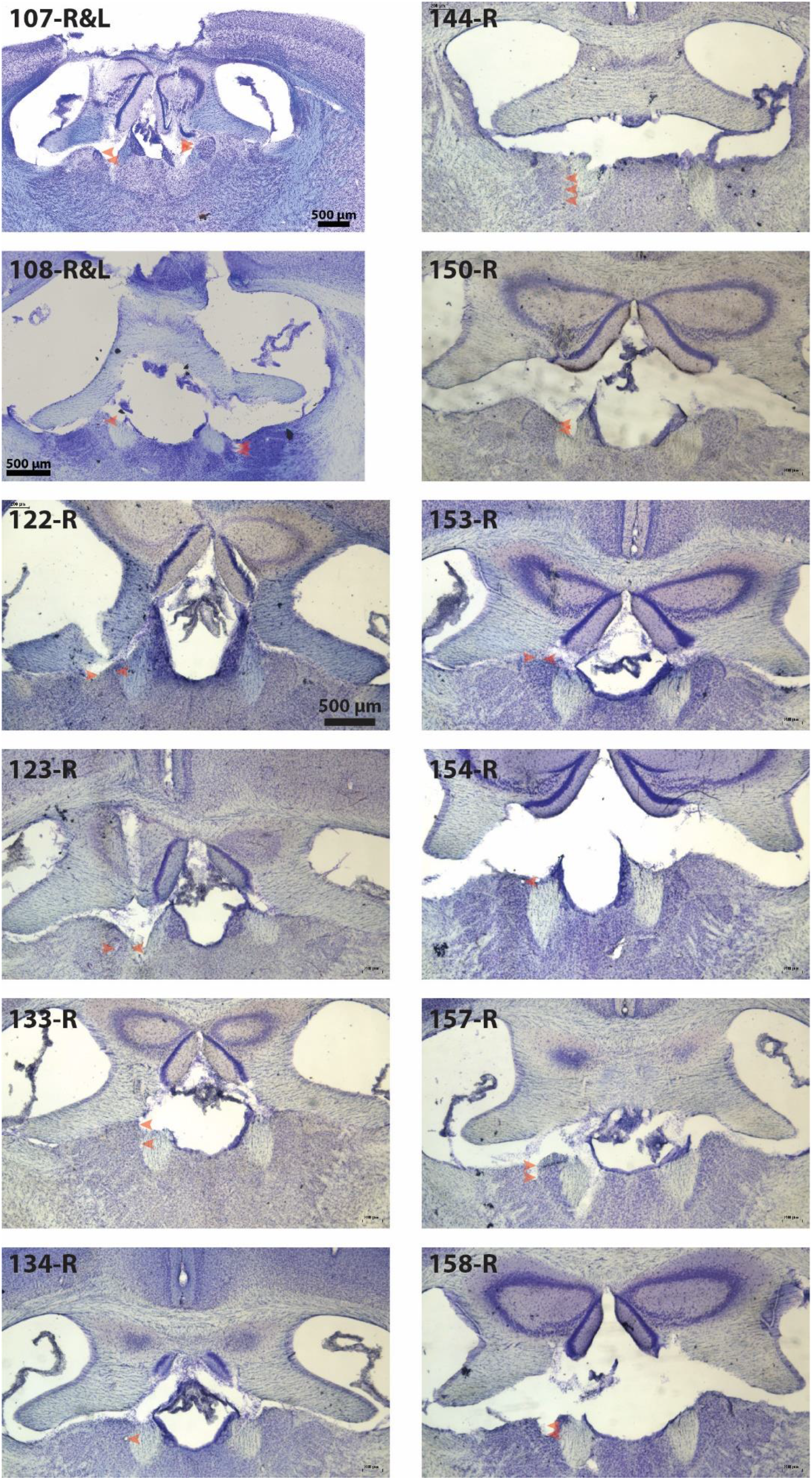
(Related to Figure 1) Histological confirmation of recording sites in the anterodorsal thalamic nucleus for L7-PKCI mice and control littermates. The scale bar provided for mouse number 122 is valid for mouse number 123-158.

**Figure S2.**
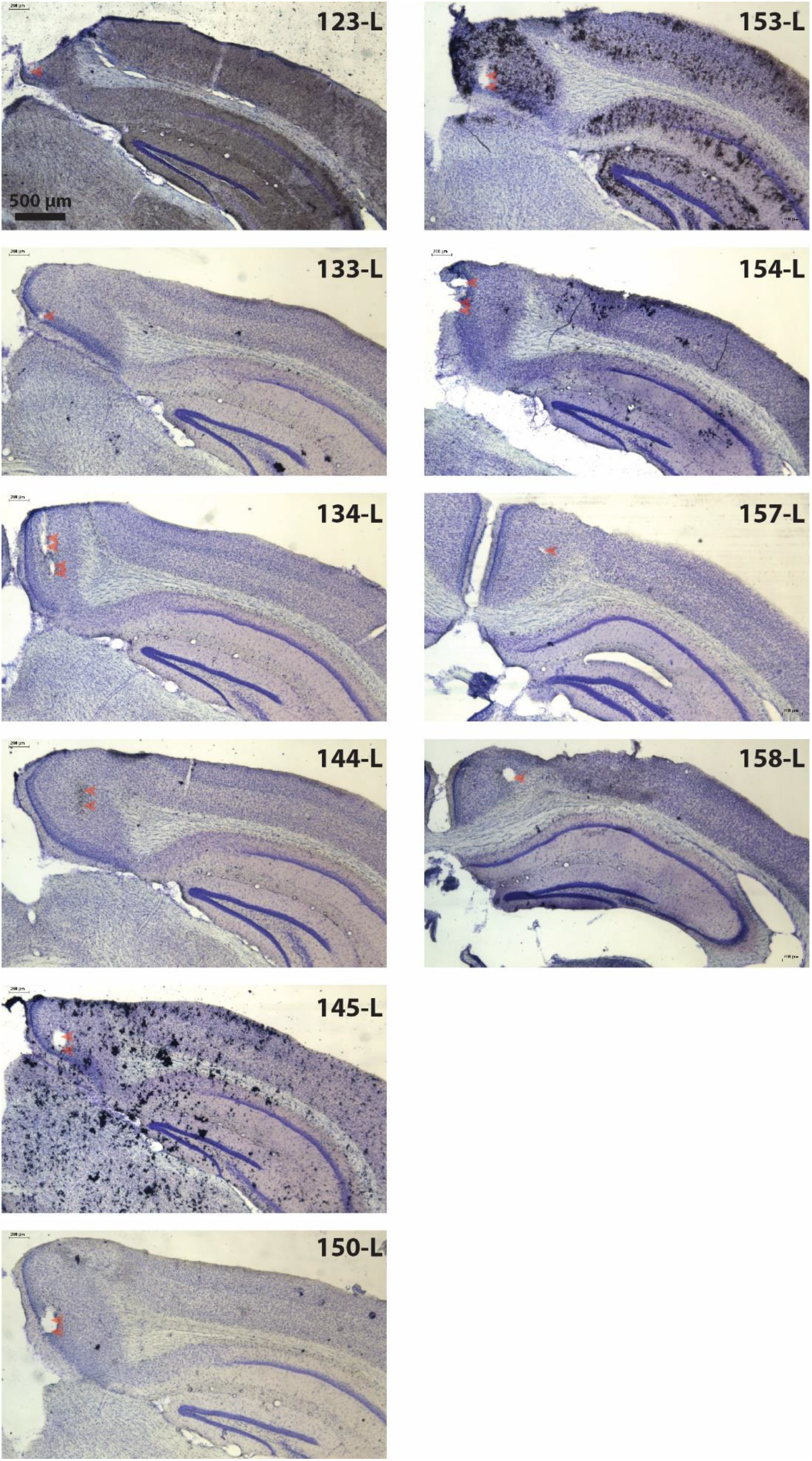
(Related to Figure 1) Histological confirmation of recording sites in the retrosplenial cortex for L7-PKCI mice and control littermates. The scale bar provided for mouse number 123 is valid for the rest of the mice.

**Figure S3.**
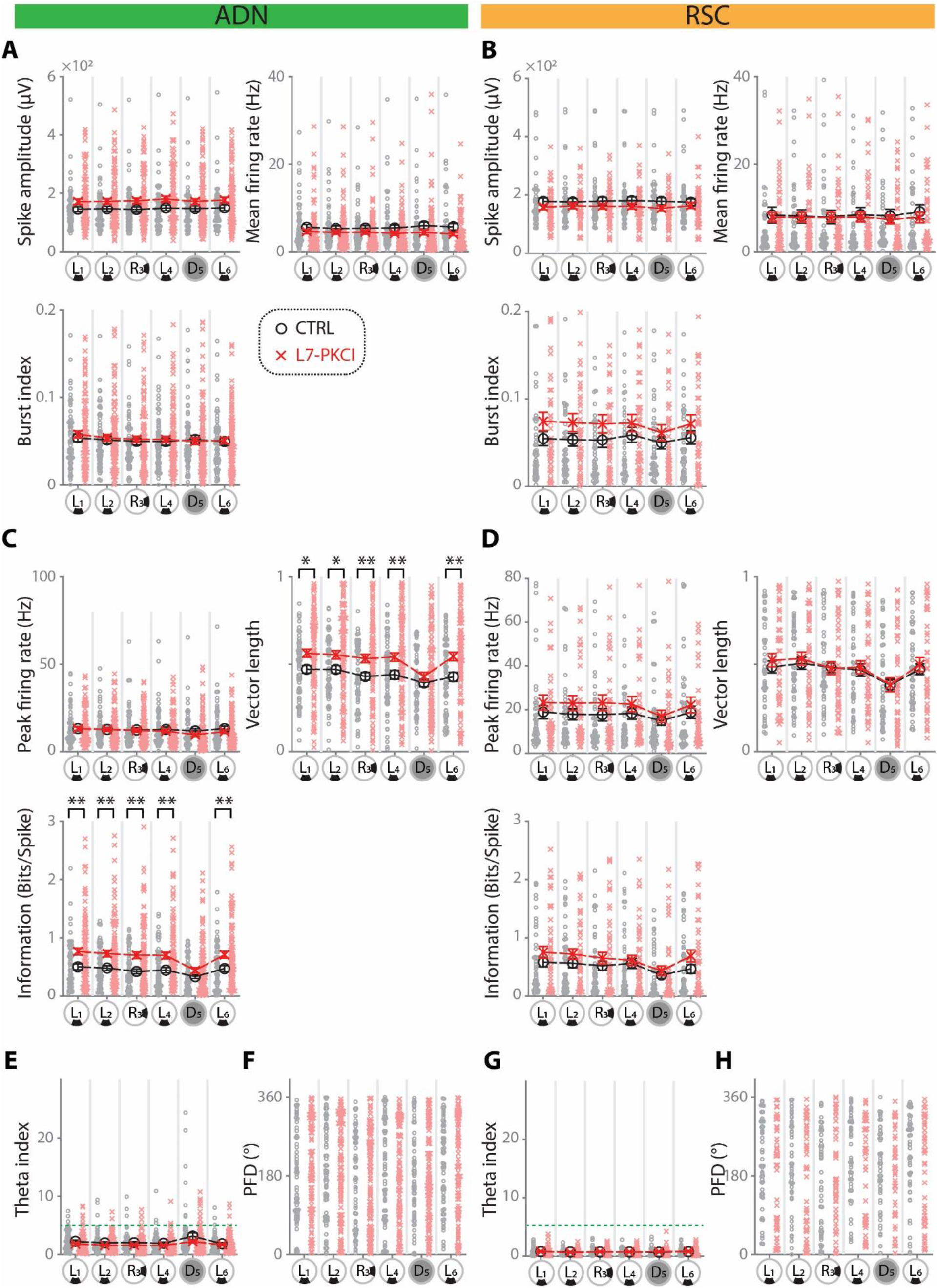
(Related to Figure 1) Basic Firing Properties and Directional Firing Properties of HD Cells in the L7-PKCI Mice. (A, B) Basic firing properties represented as spike amplitude (ADN: group × session, F_3.4, 538.5_ = 0.88, p = 0.46 two-way ANOVA; RSC: group × session, F_4.9, 443.6_ = 2.64, p = 0.023 two-way ANOVA, p > 0.1 for all session comparisons, post-hoc Bonferroni), mean firing rate (ADN: group × session, F_3.2, 504_ = 1.30, p = 0.27 two-way ANOVA; RSC: group × session, F_3.0, 268.8_= 0.92, p = 0.43 two-way ANOVA) and Burst index (ADN: group × session, F_4.2, 656.4_ = 0.94, p = 0.44 two-way ANOVA; RSC: group × session, F_3.6, 322.4_= 1.51, p = 0.21 two-way ANOVA) of HD cells from L7-PKCI mice were indifferent from that of control. (C, D) HD firing properties represented as peak firing rate (derived from the HD rate map), vector length (derived as the length of resultant vector from HD rate map), Shannon information measure, and concentration factor kappa (derived from fitting a von Mises distribution to the HD rate map). Peak firing rate was indifferent between groups (ADN: group × session, F_3.4, 529.4_ = 2.08, p = 0.09 two-way ANOVA; RSC: group × session, F_3.3, 297.3_ = 1.52, p = 0.21 two-way ANOVA). The L7-PKCI mice HD cells from ADN (C) but not RSC(D) in the light sessions showed significantly higher vector length (ADN: group × session, F_3.5, 545.4_ = 2.9, p = 0.027 two-way ANOVA, *p<0.05, **p<0.01, post-hoc Bonferroni; RSC: group × session, F_3.5, 319.0_ = 0.32, p = 0.85 two-way ANOVA), and higher information contents (ADN: : group × session, F_3.7, 574.8_ = 4.2, p = 0.003 two-way ANOVA, **p<0.01, post-hoc Bonferroni; RSC: group × session, F_4.4, 399.2_ = 1.69, p = 0.14 two-way ANOVA) compared to control. (E, G) Distribution of preferred firing direction (PFD) in HD cells of L7-PKCI from ADN (C) and RSC (G) were indifferent from control (p>0.05 for all six sessions, Watson-Williams test, Bonferroni corrected). (F, H) Theta index of HD cells from L7-PKCI mice were indifferent from that of control (ADN: group × session, F_2.6, 406.2_ = 1.58, p = 0.19 two-way ANOVA; RSC: group × session, F_4.6, 413.3_ = 0.38, p = 0.85 two-way ANOVA). Green dashed lines represent theta index of 5, values above which are considered theta modulated cells.

**Figure S4.**
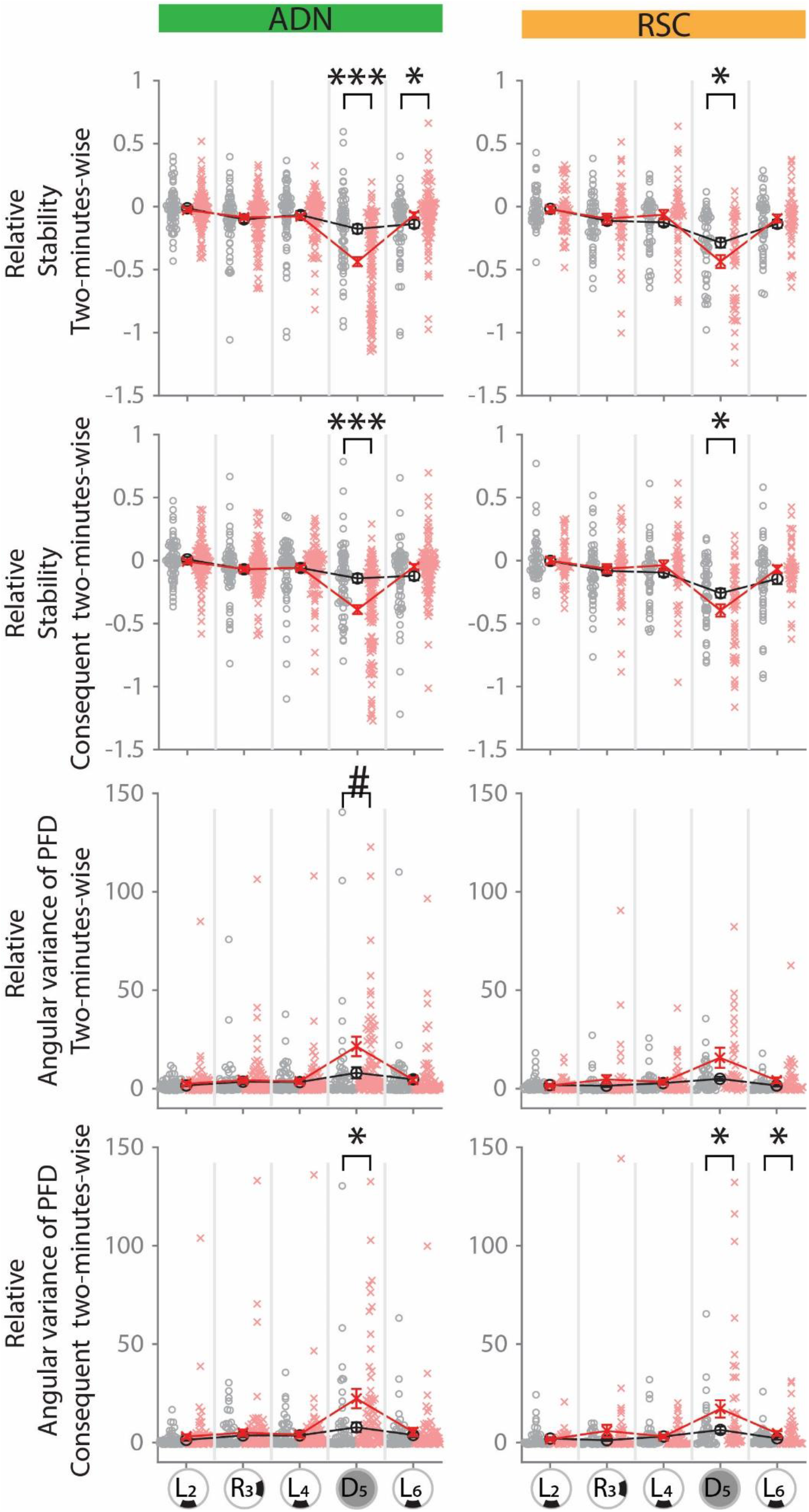
(Related to Figure 1) Complementary Analysis Showing Unstable HD Coding in L7-PKCI Mice when Relying on Self-Motion Cues. (A) Mean Pearson correlation derived from comparing all pairs of five two-minute-long HD rate maps within each 10-minutes session for ADN (Left, group × session, F_2.5, 388.6_ = 15.9, p < 0.001 two-way ANOVA; ***p < 0.001 *p < 0.05, post-hoc Bonferroni) and RSC (Right, group × session, F_3.0, 267.1_ = 4.2, p < 0.01 two-way ANOVA; *p < 0.05, post-hoc Bonferroni). (B) Mean Pearson correlation derived from comparing consecutive pairs of five two-minute-long HD rate maps within each 10-minutes session for ADN (Left, group × session, F_2.5, 397.9_ = 14.7, p < 0.001 two-way ANOVA; ***p < 0.001, post-hoc Bonferroni) and RSC (Right, group × session, F_3.2, 287.5_ = 3.9, p < 0.01 two-way ANOVA; *p < 0.05, post-hoc Bonferroni). (C) Mean angular variance of PFD derived from combining all pairs of five two-minute-long HD rate maps within each 10-minutes session for ADN (Left, group × session, F_1.3, 198.4_ = 3.8, p = 0.042 two-way ANOVA; #p = 0.053, post-hoc Bonferroni) and RSC (Right, group × session, F_1.2, 11.9_ = 3.0, p = 0.078 two-way ANOVA). (D) Mean angular variance of PFD derived from combining consecutive pairs of five two-minute-long HD rate maps within each 10-minutes session for ADN (Left, group × session, F_1.4, 222.5_ = 4.2, p = 0.029 two-way ANOVA; *p < 0.05, post-hoc Bonferroni) and RSC (Right, group × session, F_1.3,119.6_ = 4.1, p = 0.033 two-way ANOVA; *p < 0.05, post-hoc Bonferroni).

**Figure S5.**
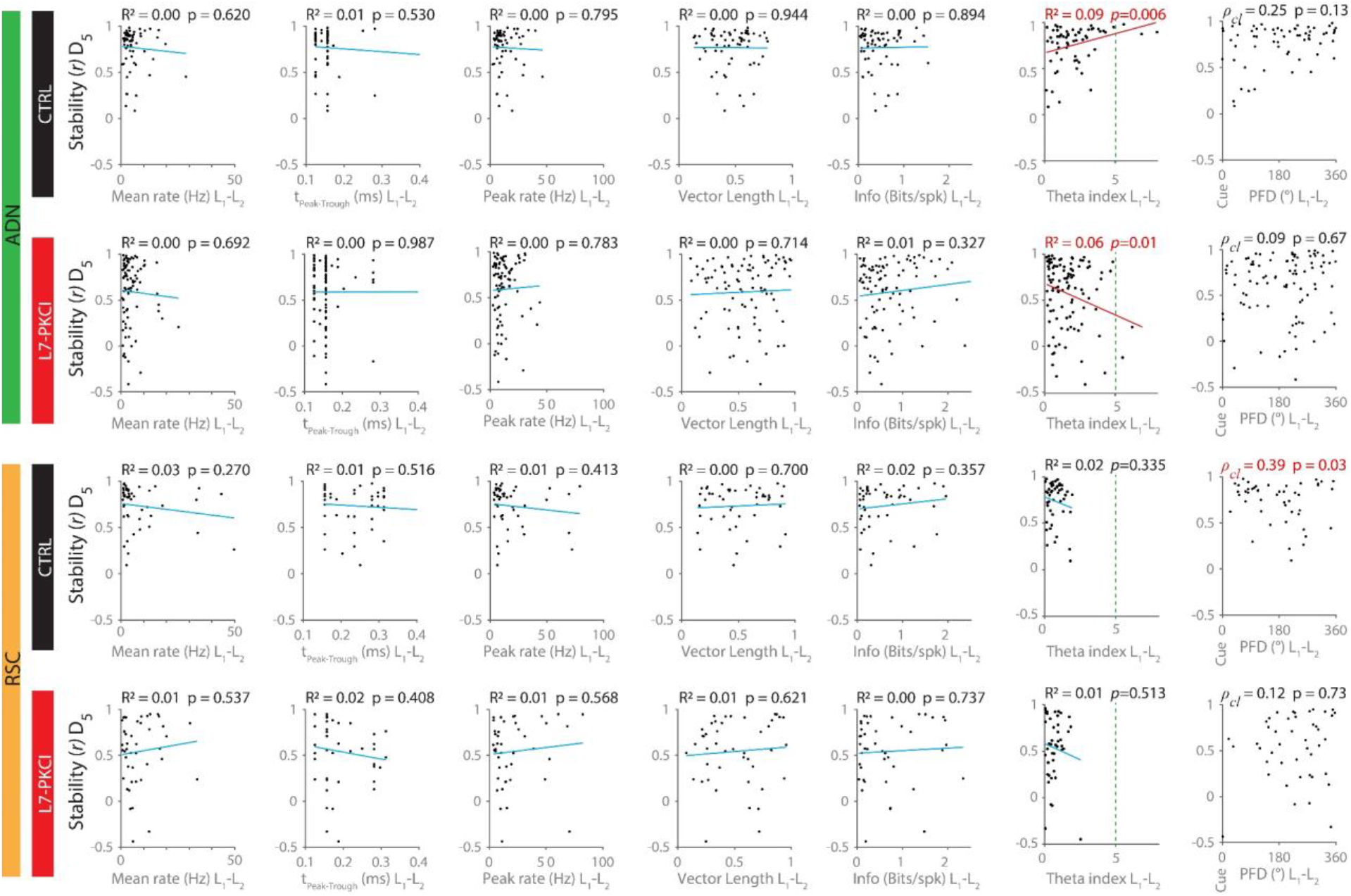
(Related to Figure 1) Instability of HD Cells in L7-PKCI under the Dark Could Not Be Assigned to any Subclass of HD Cells. The correlations between HD cells’ stability within the dark and the HD cells’ firing properties were assessed to identify whether the effect observed on the dark session is related to any HD cell subclass. Mean scores of each firing parameter from the first two standard light sessions were assigned as the score for that parameter. Linear regression models were fitted by setting HD firing stability value within the dark session as the dependent variable with each of those parameters as an explanatory (independent) variable. The parameters used included: basic firing property variables (A, mean rate; B, spike time-width), an HD specificity variable (C, Peak rate; D, Vector length; E, Shannon Information; F, concentration factor ĸ; H PFD} or a theta index (G, green dash line rep) were used as the covariates. None of these models were significant for L7-PKCI HD cells except for theta index in the ADN group (G). In the ADN group, Although very few HD cells from both control (3 HD cells) and L7-PKCI (2 HD cells) mice could be identified as a theta-modulated cell (theta index > 5), the overall stability within dark in both groups was correlated with theta index. Meanwhile, while HD cell stability within the dark and theta index were positively correlated in control, a reverse correlation was observed in L7-PKCI mice. In the RSC group, there were no theta-modulated HD cells in both control and L7-PKCI mice, and no correlations between HD cell stability within the dark and theta index of HD cell firing were observed in either of groups.

**Figure S6.**
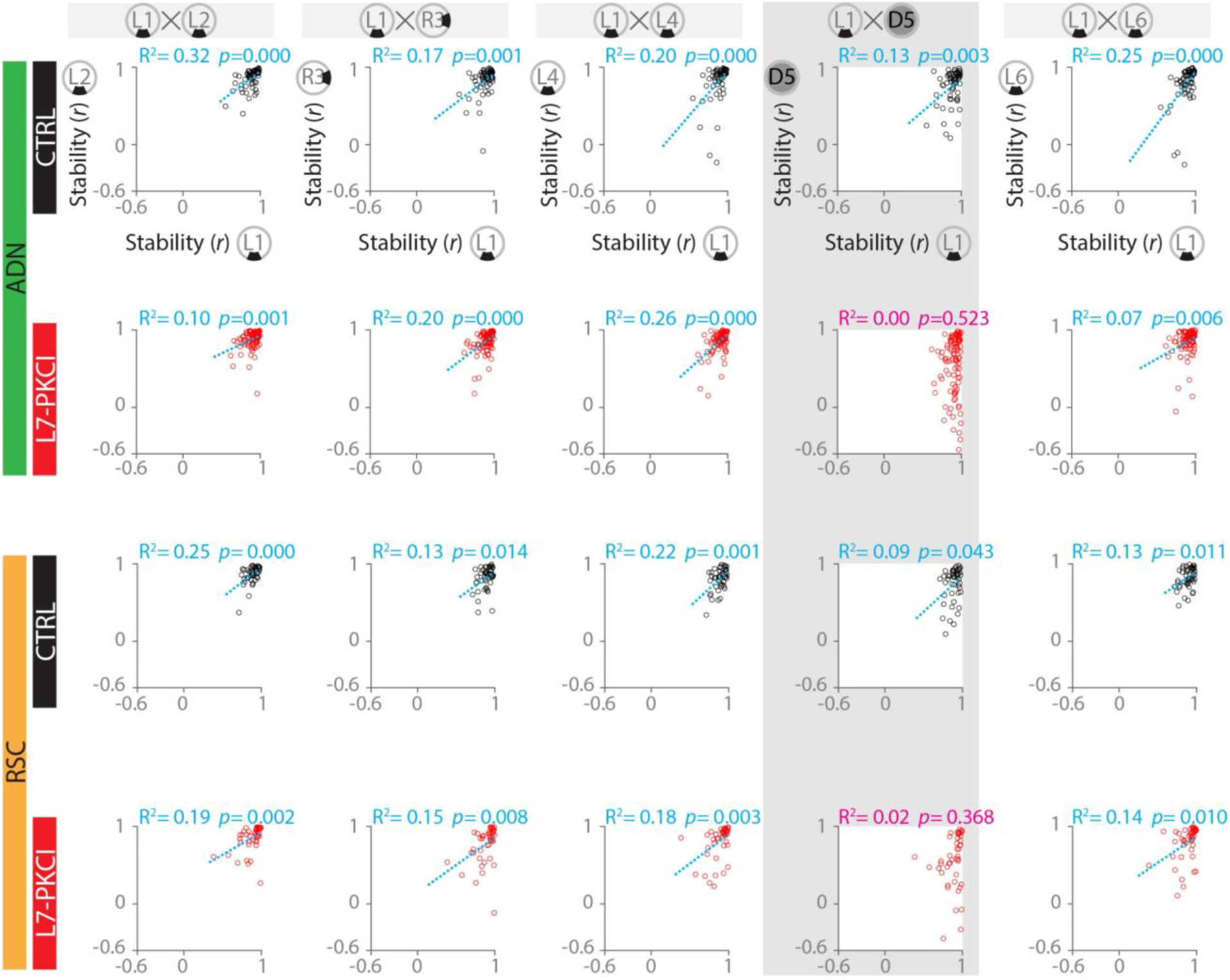
(Related to Figure 1) Correlated Inter-Session Stability of HD Rate maps is abolished in L7-PKCI Mice when Relying on Self-Motion Cues. The correlation between the stability of cells within any session with sessions L1 was assessed. HD cells of both ADN and RSC from control mice show a correlation between stability measures across all sessions. However, although HD cells of both ADN and RSC from L7-PKCI mice show a correlation in their stability measure across all sessions under the light, the correlation is abolished within the dark session.

**Figure S7.**
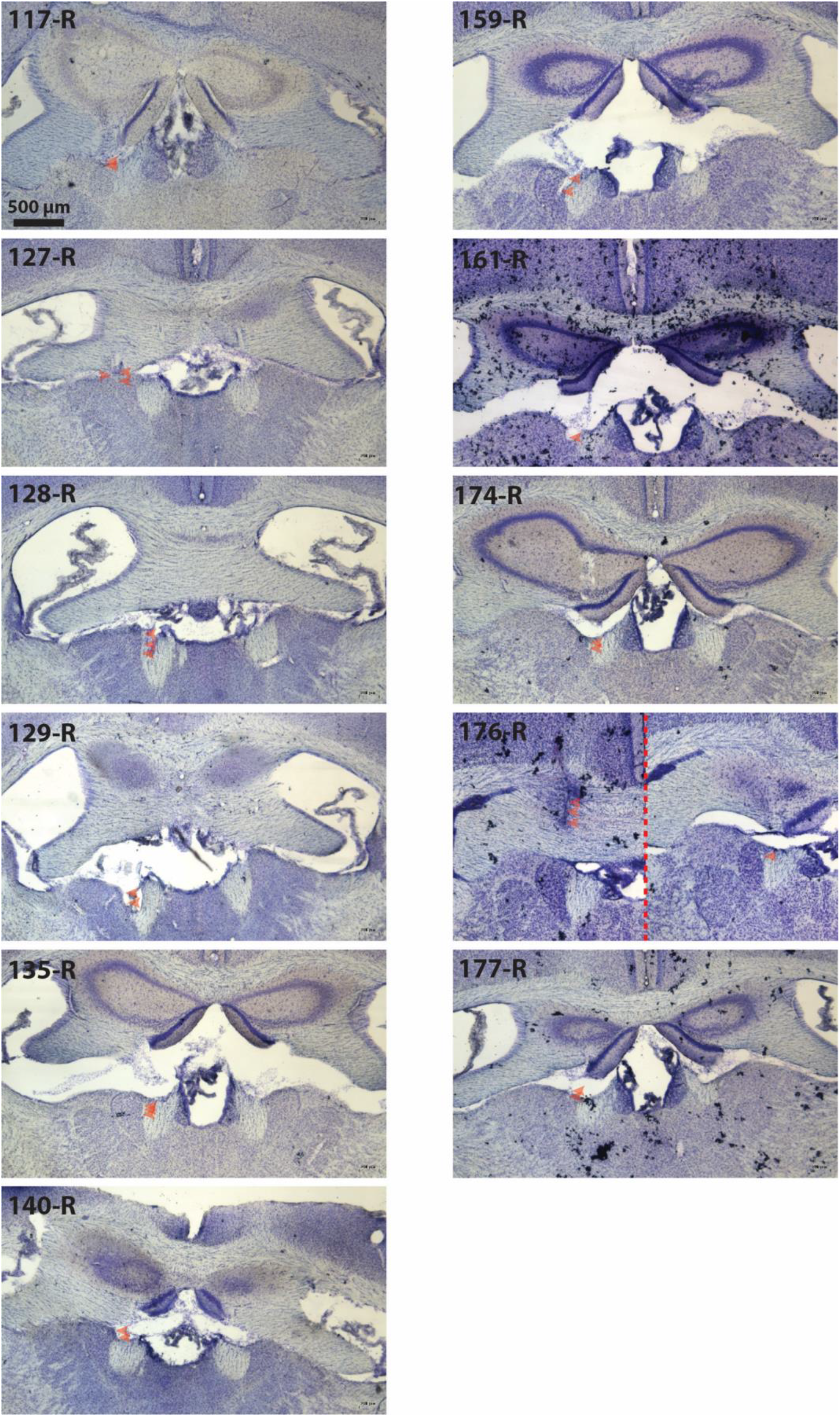
(Related to Figure 2) Histological confirmation of recording sites in the anterodorsal thalamic nucleus for L7-PP2B mice and control littermates. The scale bar provided for mouse number 117 is valid for the rest of the mice.

**Figure S8.**
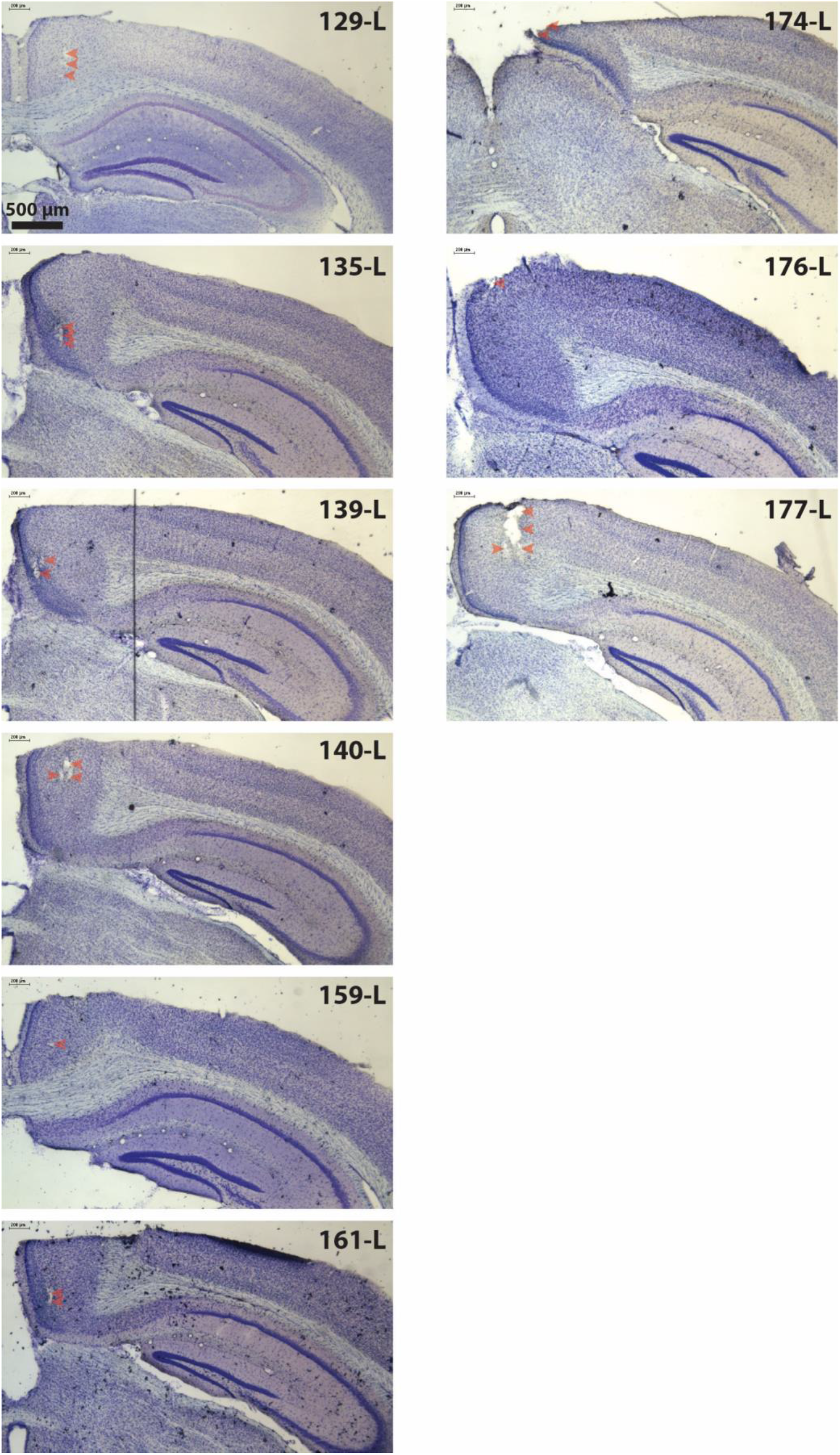
(Related to Figure 2) Histological confirmation of recording sites in the retrosplenial cortex for L7-PP2B mice and control littermates. The scale bar provided for mouse number 129 is valid for the rest of the mice.

**Figure S9.**
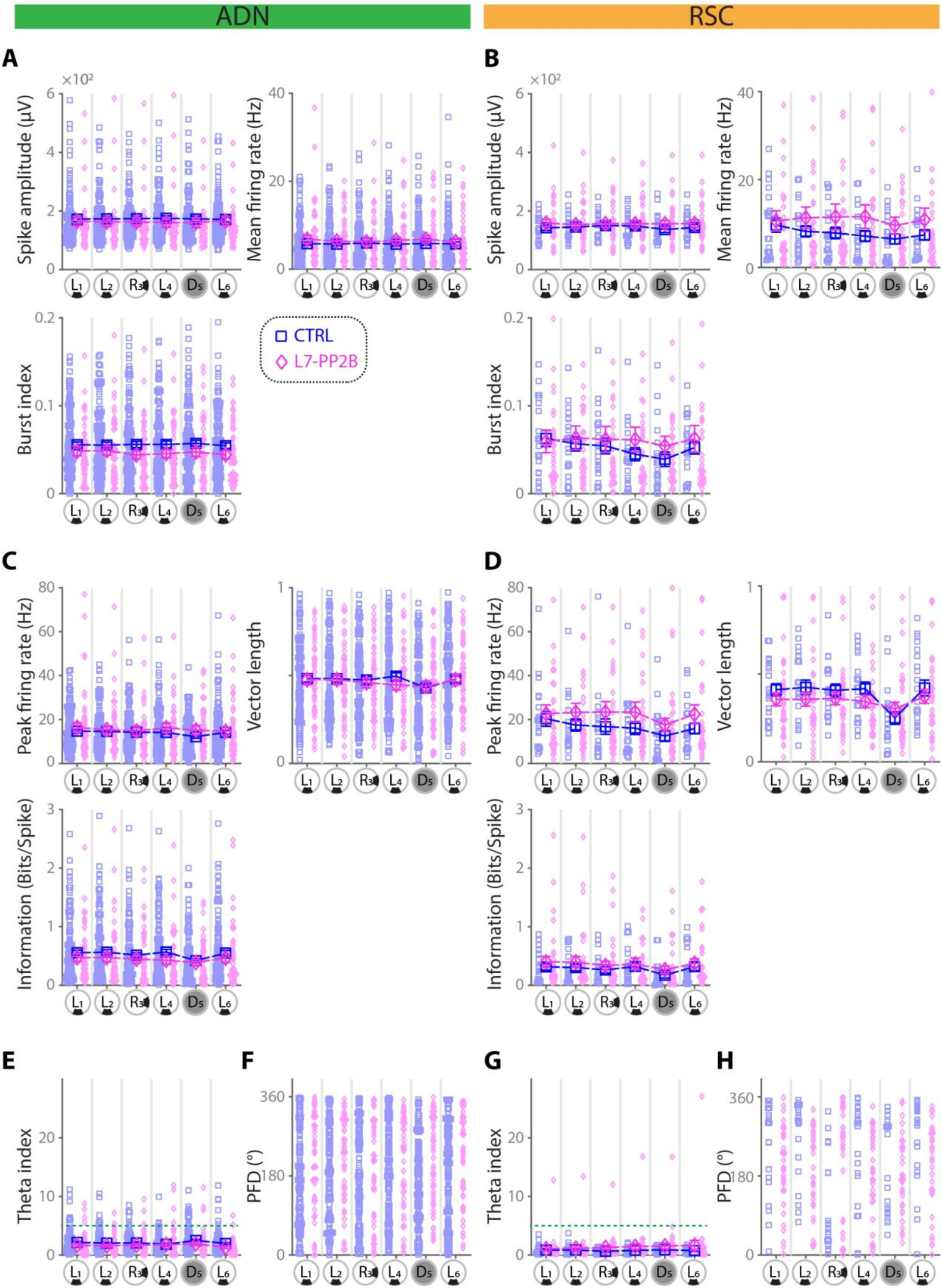
(Related to Figure 2) Basic Firing Properties and Directional Firing Properties of HD Cells in the L7-PP2B Mice. (A, B) Basic firing properties represented as spike amplitude (ADN: group × session, F_3.7, 655.4_ = 1.38, p = 0.24 two-way ANOVA; RSC: group × session, F_4.5, 223.7_ = 1.54, p = 0.18 two-way ANOVA), mean firing rate (ADN: group × session, F_3.7, 652.9_ = 0.85, p = 0.49 two-way ANOVA; RSC: group × session, F_2.6, 129.2_ = 2.35, p = 0.085 two-way ANOVA) and Burst index (ADN: group × session, F_5.0, 800.4_ = 0.57, p = 0.7 two-way ANOVA; RSC: group × session, F_2.8, 142.2_ = 2.8, p = 0.047 two-way ANOVA, p > 0.1 for all session comparisons, post-hoc Bonferroni) of HD cells from L7-PP2B mice were indifferent from that of control. (C, D) HD firing properties represented as peak firing rate (derived from the HD rate map), vector length (derived as the length of resultant vector from HD rate map), Shannon information measure, and concentration factor kappa (derived from fitting a von Mises distribution to the HD rate map). HD cells of L7-PP2B showed indifferent peak firing rate (ADN: group × session, F_3.7, 659.1_ = 2.15, p = 0.078 two-way ANOVA; RSC: : group × session, F_2.8, 139.7_ = 1.44, p = 0.23 two-way ANOVA), vector length (ADN: group × session, F_4.6, 812.3_ = 1.8, p = 0.13 two-way ANOVA; RSC: group × session, F_4.4, 219.6_ = 3.9, p = 0.003 two-way ANOVA, p > 0.1 for all session comparisons, post-hoc Bonferroni), and information contents (ADN: group × session, F_4.5, 807.4_ = 2.07, p = 0.07 two-way ANOVA; RSC: group × session, F_3.1, 154.6_ = 0.28, p = 0.84 two-way ANOVA) compared to control. (E, G) Distribution of preferred firing direction (PFD) in HD cells of L7-PP2B from ADN (E) and RSC (G) were indifferent from control (p>0.05 for all six sessions, Watson-Williams test, Bonferroni corrected). Green dashed lines represent theta index of 5, values above which are considered theta modulated cells. (F, H) Theta index of HD cells from L7-PP2B mice were indifferent from that of control (ADN: group × session, F_4.5, 798.6_ = 0.94, p = 0.45 two-way ANOVA; RSC: group × session, F_1.7, 82.9_ = 0.88, p = 0.40 two-way ANOVA).

**Figure S10.**
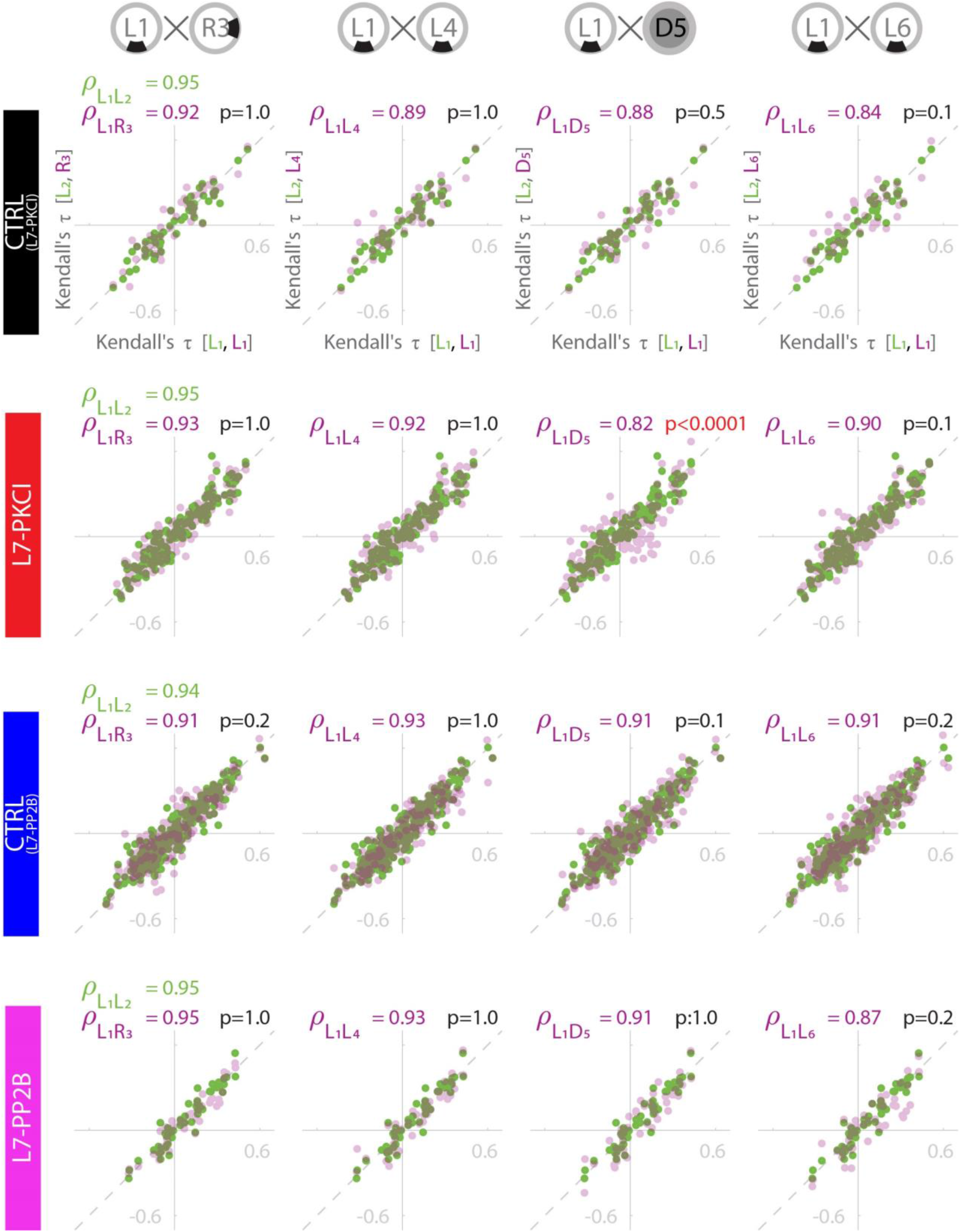
(Related to Figure 3) Complementary Analysis of Temporal Coordination Demonstrating a Disrupted Temporal Coordination of L7-PKCI HD Cells under the Dark. Pearson correlation between t_Kendall_ of each cell pair from each session with t_Kendall_ of that cell pair in session L1. Pearson correlation between session L2 and L1 (green) is represented as the baseline, and the correlation between t_Kendall_ of any other session with session L1 (Purple) is compared to this baseline. The t_Kendall_ for each simultaneously recorded pair of HD cells was measured by Kendall’s correlation between spike timestamps in a 1000-ms time window. Note that only the HD cell pairs from L7-PKCI mice and only within the dark session represent a correlation significantly lower than their baseline.

**Figure S11.**
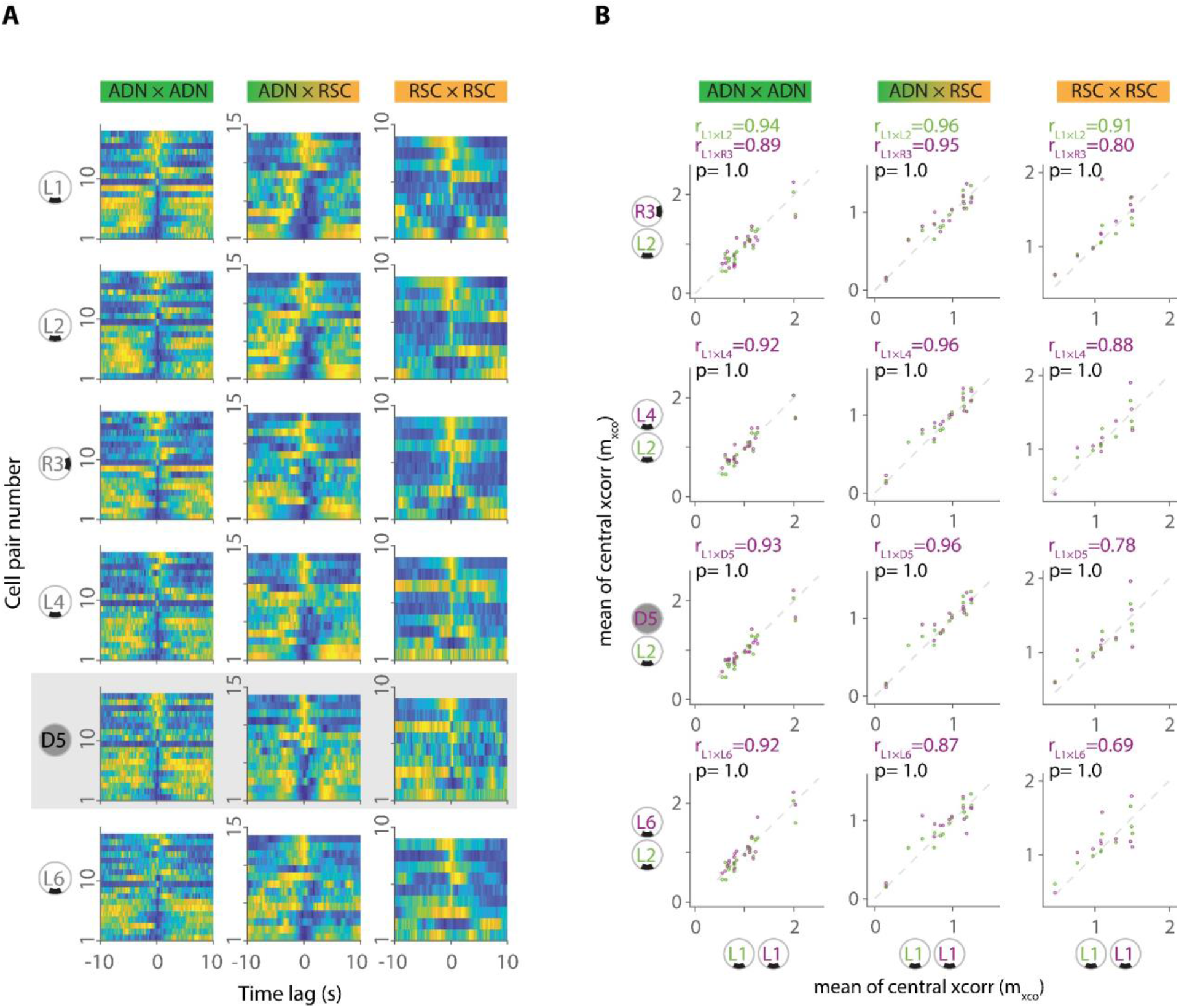
(Related to Figure 4) Intact Temporal Coordination within ADN, within RSC, and between ADN and RSC HD Cells of L7-PP2B Mice. (A) Temporal cross-correlograms of all simultaneously recorded HD cell pairs in L7-PP2B mice within ADN (left), between ADN and RSC (middle), and within RSC (right) across all sessions. Within each heat-map, each row represents the cross-correlogram of one HD cell pair. (B) Pearson correlation between m_xco_ of each session and session L1 indicates that the correlation structure between ADN and RSC cells (Middle) and within each of ADN (Top) and RSC (Bottom) has remained intact. In each comparison, the correlation between session L2 and L1 (green) is used as the baseline, and the correlation between m_xco_ of any other session and session L1 (purple) is compared to this baseline. The inset value in each scatter plot represents the correlation value between L1 and L2 (Top, green), the correlation value between L1 and the session indicated in purple (Middle purple), and the p-value from comparing the two correlations (green versus purple distribution). The p-Values are corrected after the Bonferroni approach for multiple comparisons.

**Figure S12.**
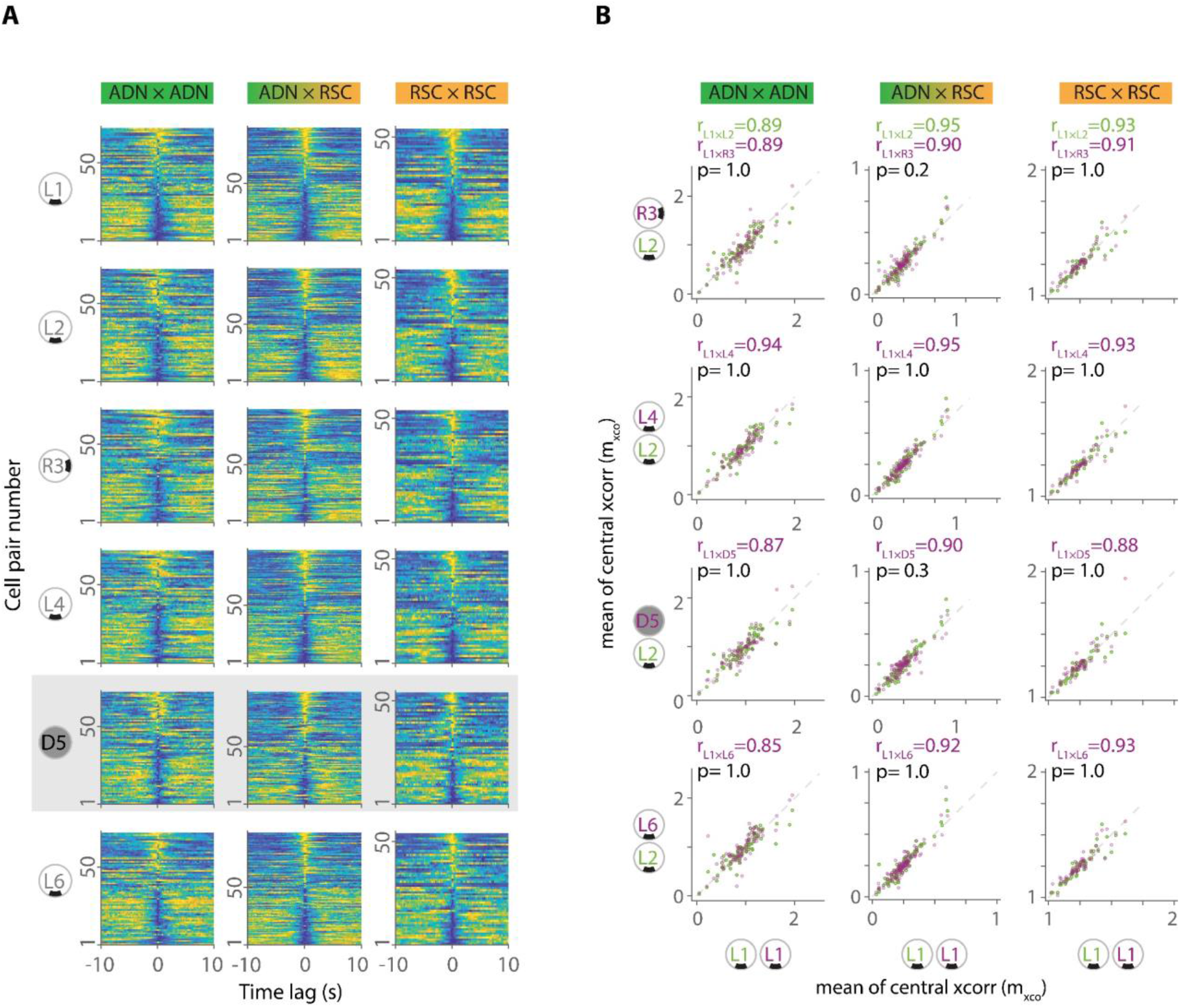
(Related to Figure 4) Intact Temporal Coordination within ADN, within RSC, and between ADN and RSC HD Cells of L7-PP2B Mice. (A) Temporal cross-correlograms of all simultaneously recorded HD cell pairs in L7-PP2B mice within ADN (left), between ADN and RSC (middle), and within RSC (right) across all sessions. Within each heat-map, each row represents the cross-correlogram of one HD cell pair. (B) Pearson correlation between m_xco_ of each session and session L1 indicates that the correlation structure between ADN and RSC cells (Middle) and within each of ADN (Top) and RSC (Bottom) has remained intact. In each comparison, the correlation between session L2 and L1 (green) is used as the baseline, and the correlation between m_xco_ of any other session and session L1 (purple) is compared to this baseline. The inset value in each scatter plot represents the correlation value between L1 and L2 (Top, green), the correlation value between L1 and the session indicated in purple (Middle purple), and the p-value from comparing the two correlations (green versus purple distribution). The p-Values are corrected after the Bonferroni approach for multiple comparisons.

**Figure S13.**
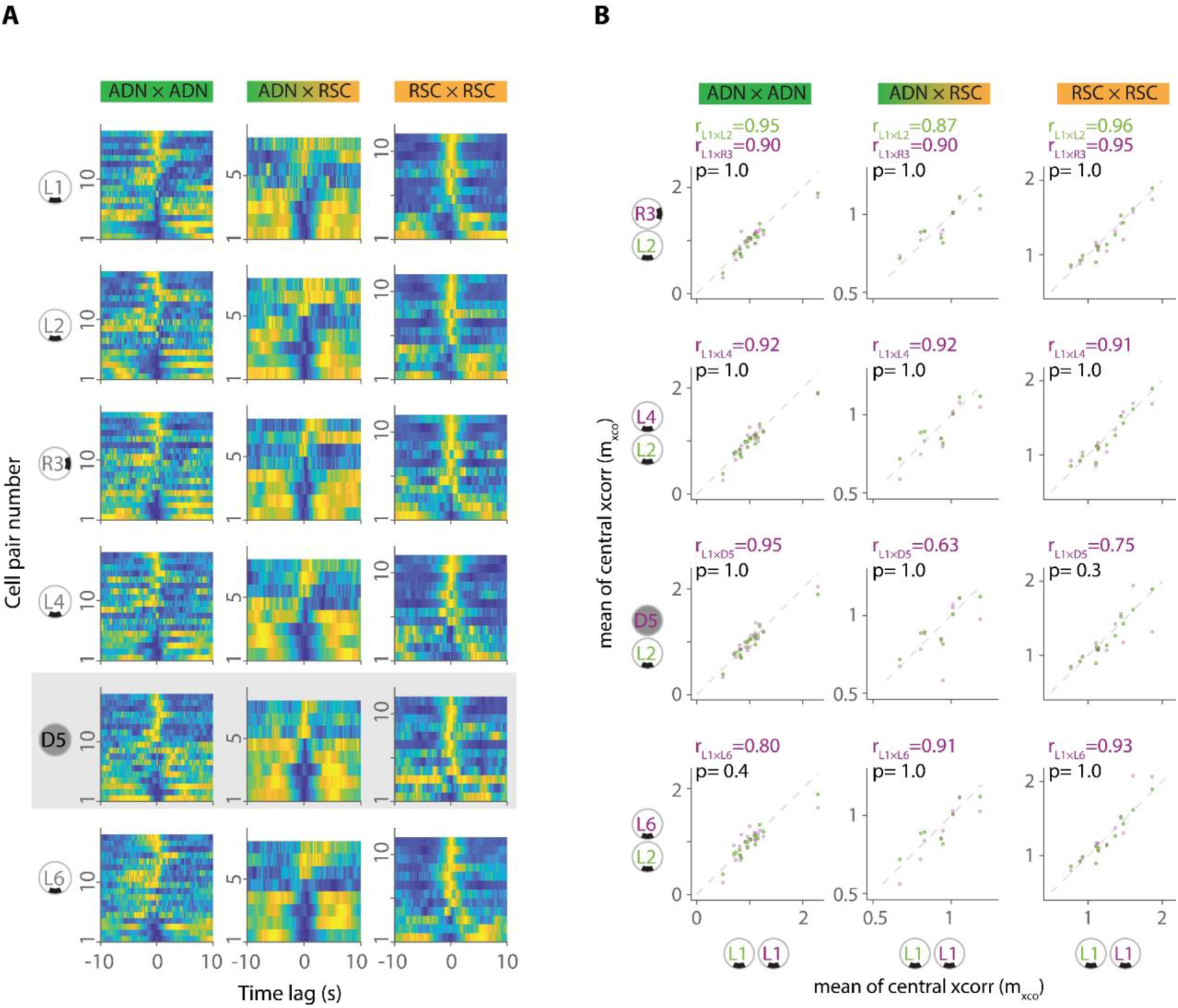
(Related to Figure 4) Intact Temporal Coordination within ADN, within RSC, and between ADN and RSC HD Cells in control littermates of L7-PP2B Mice. (A) Temporal cross-correlograms of all simultaneously recorded HD cell pairs in control littermates of L7-PP2B mice within ADN (left), between ADN and RSC (middle), and within RSC (right) across all sessions. Within each heat-map, each row represents the cross-correlogram of one HD cell pair. (B) Pearson correlation between m_xco_ of each session and session L1 indicates that the correlation structure between ADN and RSC cells (Middle) and within each of ADN (Top) and RSC (Bottom) has remained intact. In each comparison, the correlation between session L2 and L1 (green) is used as the baseline, and the correlation between m_xco_ of any other session and session L1 (purple) is compared to this baseline. The inset value in each scatter plot represents the correlation value between L1 and L2 (Top, green), the correlation value between L1 and the session indicated in purple (Middle purple), and the p-value from comparing the two correlations (green versus purple distribution). The p-Values are corrected after the Bonferroni approach for multiple comparisons.

**Figure S14.**
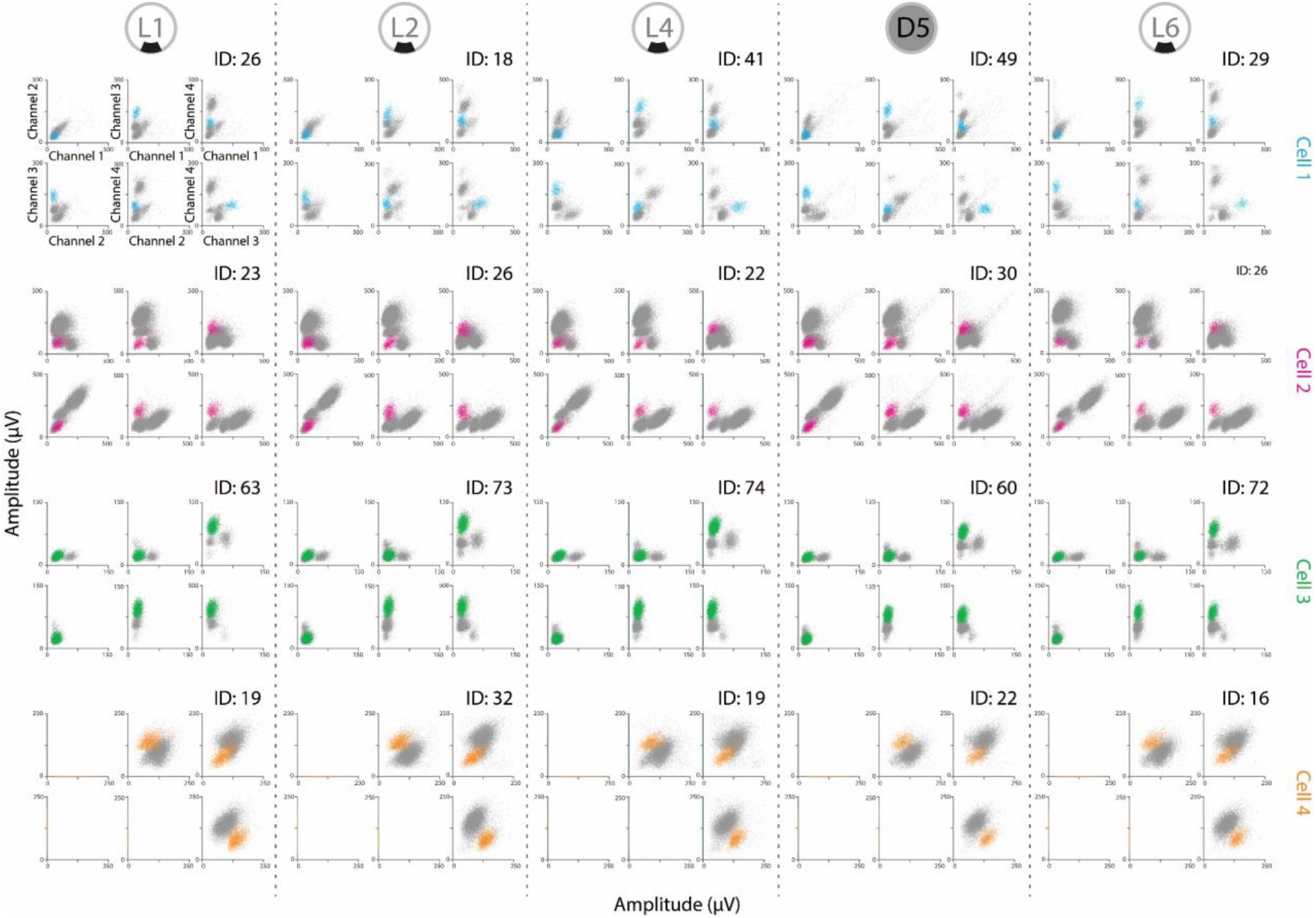
(Related to Figure 4) Cluster Stability and Cluster Quality of the Four Representative HD Cells Used in. Figure 4 Cluster quality of each cell (colored dots) within each session respective to the rest of the spikes (gray dots) is estimated by an isolation distance measure (ID; Supplemental Experimental Procedures) which are given on the top-right of each session. Note the stability of each cluster across all sessions and their ID values above 10 in all sessions.

**Figure S15.**
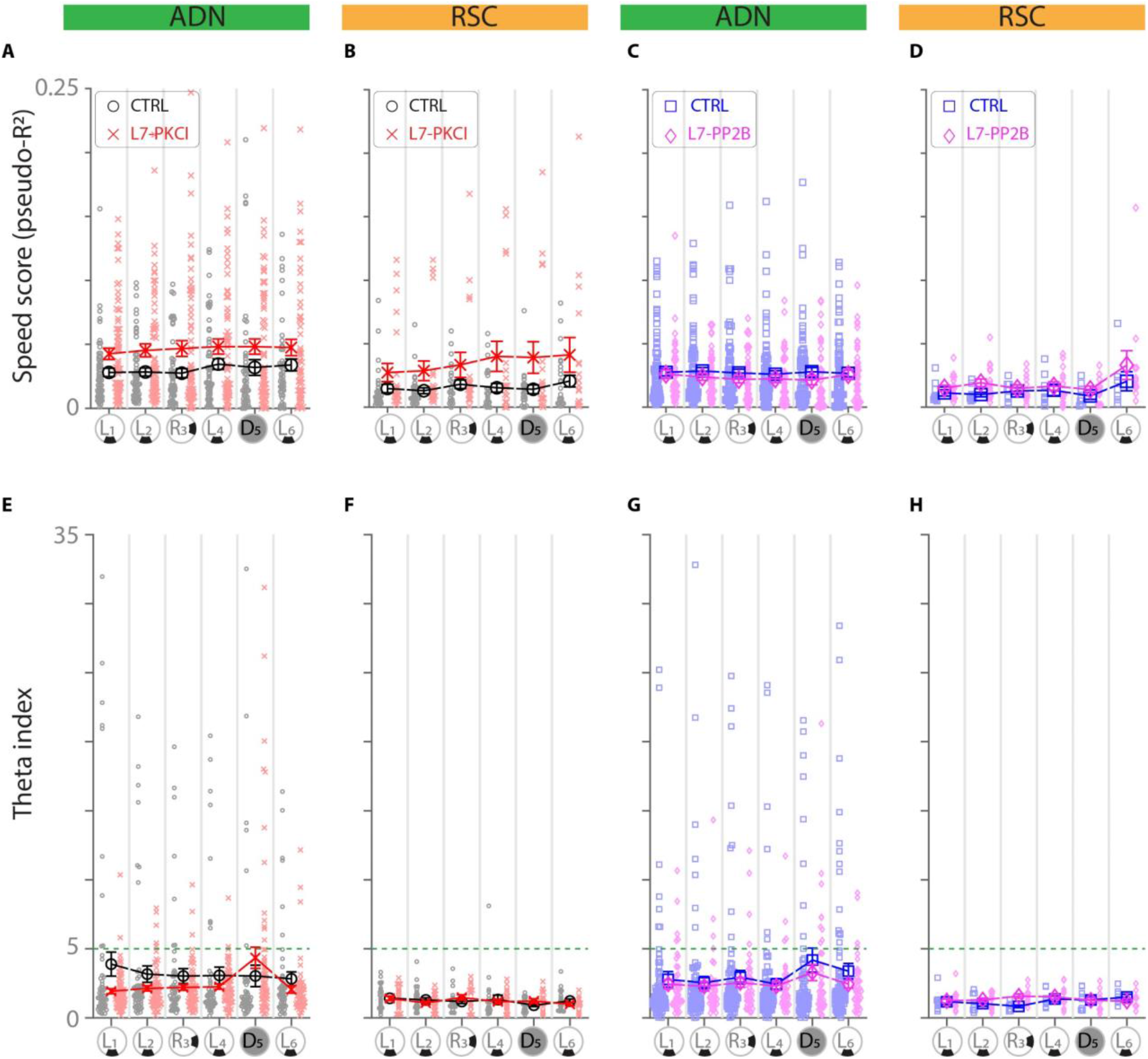
(Related to Figures 5 and 6) Complementary Analysis on Speed Cells. (A-B) Speed cells in L7-PKCI mice exhibited an overall stronger speed scores compared to control in both ADN (A, group main effect, p = 0.013) and RSC (B, group main effect, p = 0.032), while no interaction effect was observed (ADN: group × session, F_3.3, 306.7_ = 0.79, p = 0.51, two-way ANOVA; RSC: group × session, F_3.1, 85.7_ = 1.10, p = 0.35, two-way ANOVA). (C-D) Speed cells in L7-PP2B mice exhibited indistinguishable speed scores compared to control in both ADN (C, group main effect, p = 0.21, and group × session, F_3.5, 377.4_ = 0.53, p = 0.69, two-way ANOVA) and RSC (D, group main effect, p = 0.52, and group × session, F_1.2, 16.8_ = 0.31, p = 0.63, two-way ANOVA). (E-F) Theta index of speed cells in all four groups was similar to their respective controls.

**Table S1.** List of mice and their groupings with the types of cells recorded from each mouse.

| MOUSE# | GROUP | RIGHT HEM (R) |  |  |  | LEFT HEM (L) |  |  |  |
| --- | --- | --- | --- | --- | --- | --- | --- | --- | --- |
|  |  | Target | HD | AHV | Speed | Target | HD | AHV | Speed |
| 107 | PKCI-WT | ADN | 1 | N | Y | ADN | Y | N | Y |
| 108 | PKCI-MU | ADN | 1 |  |  | ADN | Y | Y | Y |
| 122 | PKCI-WT | ADN | Y | N | Y | RSC | N | N | N |
| 123 | PKCI-MU | ADN | Y | N | Y | RSC | Y | N | Y |
| 133 | PKCI-MU | ADN | Y | N | Y | RSC | Y | Y | Y |
| 134 | PKCI-WT | ADN | Y | Y | Y | RSC | Y | Y | Y |
| 144 | PKCI-WT | ADN | Y | Y | Y | RSC | Y | Y | Y |
| 145 | PKCI-MU | ADN | 1 | N | Y | RSC | N | 1 | N |
| 150 | PKCI-WT | ADN | 1 | N | Y | RSC | Y | N | Y |
| 153 | PKCI-WT | ADN | Y | 1 | Y | RSC | Y | Y | Y |
| 154 | PKCI-MU | ADN | Y | Y | Y | RSC | Y | 1 | Y |
| 157 | PKCI-WT | ADN | Y | Y | Y | RSC | Y | Y | Y |
| 158 | PKCI-MU | ADN | Y | N | Y | RSC | Y | 1 | Y |
|  |  | Target | HD | AHV | Speed | Target | HD | AHV | Speed |
| 117 | PP2B-MU | ADN | 1 | Y | Y | RSC | N | N | N |
| 127 | PP2B-WT | ADN | Y | N | Y | RSC | N | N | N |
| 128 | PP2B-WT | ADN | Y | N | Y | RSC | Y | Y | Y |
| 129 | PP2B-MU | ADN | Y | N | Y | RSC | Y | N | Y |
| 135 | PP2B-MU | ADN | N | N | N | RSC | Y | N | Y |
| 139 | PP2B-WT | ADN | Y | N | Y | RSC | Y | N | Y |
| 140 | PP2B-MU | ADN | Y | N | Y | RSC | Y | Y | Y |
| 159 | PP2B-WT | ADN | Y | N | Y | RSC | Y | Y | Y |
| 161 | PP2B-MU | ADN | Y | N | Y | RSC | Y | Y | Y |
| 174 | PP2B-MU | ADN | Y | N | Y | RSC | Y | Y | Y |
| 176 | PP2B-WT | ADN | Y | N | Y | RSC | Y | Y | Y |
| 177 | PP2B-WT | ADN | Y | N | Y | RSC | Y | N | Y |

## EXPERIMENTAL MODEL AND SUBJECT DETAILS

### Subjects

Data were obtained from 25 adult male mice (six L7-PKCI with seven control littermates; six L7-PP2B with six control littermates), aged from three to six months during the recordings. Mice were housed singly in a standard transparent polycarbonate cage (25x16x14cm; LxWxH) on a 12-hour light/dark cycle. All the recordings were performed during the light cycle. Animals were food-restricted during recording days (minimum 85% of normal body weight measured during ad libitum diet) to increase their foraging performance. The experimenters were blind in respect to animal genotype. All experiments were performed in accordance with the official European guidelines for the care and use of laboratory animals (86/609/EEC) and in accordance with the Policies of the French Committee of Ethics (Decrees n ° 87-848 and n ° 2001-464). In the L7-PKCI mouse line, the pseudosubstrate PKC inhibitor (PKCI) is selectively expressed in Purkinje cells under the control of the pcp-2 (L7) gene promoter. L7-PKCI mice display normal cerebellar histology and an intact induction of long-term potentiation (LTP) but an impaired long-term depression (LTD) at cerebellar parallel fiber-Purkinje cell (PF-PC) synapses (Goossens et al., 2001; De Zeeuw et al., 1998). Control mice for L7-PKCI groups were their wild-type littermates. In L7-PP2B mice, the selective deletion of PP2B in Purkinje cells was obtained using the Cre-loxP-system (flox/flox, cre/+). L7-PP2B mice display normal cerebellar histology and an intact induction of LTD but an impaired induction of LTP at the PF-PC synapses. L7-PP2B mice also display defects in the baseline excitability and intrinsic plasticity of Purkinje cells. Control mice for L7-PP2B groups were their (flox/flox, +/+) littermates (Schonewille et al., 2010).

## METHOD DETAILS

### Surgery

Mice were handled and introduced to the recording apparatus to experience few foraging sessions daily for 1-2 weeks before surgery. Only pairs of littermates with satisfactory foraging performance were subjected to surgery. During surgery, mice were anesthetized deeply using isoflurane and fixed in a stereotaxic instrument. An incision was made on the skin to expose the skull. Small craniotomies on the skull were made above the anterodorsal thalamic nucleus (ADN) and/or the retrosplenial cortex (RSC). Two custom-made 3D-printed single-screw micro-drives each carrying either four (in 22 mice) or eight (in three mice) tetrodes were used for targeting either bilateral ADN (-0.5mm AP, ±0.7mm ML, implanted at 1.7 to 2.0mm ventral to brain surface) or right ADN with left RSC (-2.8 to -3.0mm AP, 0.55mm ML, implanted at 0.1 to 0.6mm ventral to brain surface). During electrophysiological recordings, tetrodes were further lowered (at the ventral direction) to reach the desired structure. Tetrodes were constructed by twisting together four strands of 17-µm polyamide-coated platinum-iridium (90%-10%) wire (California Fine Wire, USA) and were heated to melt the insulation and create stiff and closely arranged tips (recording sites). Tetrodes were housed in a 23 AWG (for 4-tetrodes micro-drive) or a 19 AWG (for 8-tetrodes micro-drive) stainless steel hypodermic tubing and assembled on a micro-drive. Tetrode wire strands were connected to a small custom-made printed circuit board (PCB) with built-in omnetics connectors (Omnetics Connector corp.). Tetrodes were delicately cut under a stereomicroscope using a stainless-steel scissor (14058-11, Fine Science Tools, CA) for creating flat, smooth, and shining tips. Tetrodes were immersed in a mixture of 75% polyethylene glycol (1 mg/mL in deionized water, sigma-Aldrich) in platinum black plating solution (Neuralynx Inc, USA) and were electroplated to reduce electrode impedances to 100-250 kO at 1kHz (Ferguson et al., 2009). All the combinations of pairs of tetrode wires were also tested for the absence of a short circuit between them. Electroplating was followed up on multiple intermittent days to ensure the presence of a long-lasting electroplated coating by measuring a stable impedance in the desired range. During surgery, four or five stainless-steel screws were mounted on the skull for securing the implantation. Two of these screws were mounted above the frontal cortex and served as grounds. The implant was fixed using super-bond (SunMedical, Japan) and dental cement. After the surgery, all mice were subjected to a minimum of 1-week recovery before recordings. After recovery and before initiating electrophysiological recordings, all mice were checked to ensure they have normal physical characteristics and sensory-motor reflexes such as cage movement, grooming, rearing, eye reflex, ear twitch, and whisker responses.

### Electrophysiology, Single-unit recording

After recovery, tetrodes were screened for head-direction cell activity while mice were foraging for chocolate cereal crumbs scattered on the surface of a white wooden circular open field (50 × 35 cm; diameter × height) covered with a matte-finish waterproof PVC cover. The arena was featureless except a removable cue card constructed by a black PVC sheet (21x29 cm) attached with an object with an isosceles triangular-shaped base (base size: 4.5x4.5x7cm; height: 21cm). The object was furnished with black and white matte-finish PVC 45°-slant strips and was centered along the length of the black PVC sheet. The arena was surrounded by a circular black floor-to-ceiling curtain (1.8m diameter). Four uniformly arranged lamps were located on the ceiling creating homogeneous 8-10 lux illumination. A speaker generating 80-85 dB white noise sound and an infrared camera were centered above the arena on the ceiling. A pulley system was used for counterbalancing the cable weight.

Electrical signals recorded on the tip of tetrodes were amplified by a unity-gain operational amplifier on two headstages (HS-18-LED for four-tetrodes microdrives, and HS-36-LED for eight-tetrodes micro-drives, Neuralynx Inc.) above the animal head and transferred to a 64-channels data acquisition system (Digital Lynx Combo Board, Neuralynx Inc.) using ultralight conner wires. The signal from each tetrode was differentially recorded against a tetrode with a low activity to detect low-noise cell activity. On the data acquisition system, electrical signals were band-pass filtered (600 Hz to 6 kHz). The filtered signal was passed through a preset amplitude-threshold to detect the extracellular action potentials and digitized at 32 kHz using a 24-bit analog-to-digital converter (Digital Lynx Combo Board and Cheetah software, Neuralynx Inc). Animal tracking (collection of position and head direction samples) was performed by monitoring the position of two differently sized custom-made infrared (860-940nm wavelength) light-emitting diode (LED) circuits distanced 6 to 8 cm apart and located on the headstage on top of animal’s head. LED circuits were built from a series of SMT (Surface-mount technology) LEDs (#876-8732, #871-4669, RS Components Ltd.) and SMT resistors creating light spots with desired sizes. The two LED circuits were fixed along the axis of the animal’s head, with the circuit generating a more prominent light spot located on the rear and the one generating a smaller light spot located in front. Animal position and head direction samples were reconstructed offline from LED tracks. Head direction samples were generated based on the vector pointing from the rear light spot toward the frontal. The animal tracking data was incorporated into the data acquisition system and generated 25Hz tracking samples synchronized with the electrical signals recorded on tetrodes.

During cluster screenings, when an isolated cluster (representing single-cell activity) was observed on any channel of all tetrodes, the animal was conditioned to two 10-min standard foraging sessions under standard light conditions with a 5-min resting inter-session interval. During a resting session, mice were moved to a remote resting spot (a small flower pot covered by a soft towel) with no visual access to the open field arena. Before the first session, the arena surface and walls were wiped with 20% ethanol to remove any olfactory cues from different mice. The arena was vacuumed between each two light sessions, wiped with soapy water, and dried to remove any olfactory cues. After each standard session, detected spikes were sorted into isolated clusters (representing individual neurons) using an automated spike sorting software (Snap, Neuralynx Inc., USA). The firings of cells were analyzed to detect head-direction cell activity using a custom-made program in MATLAB. If any isolated clusters exhibited HD cell activity in at least one of the two sessions, the animal was conditioned to a full-protocol experiment; otherwise, tetrodes were advanced 15-60 µm, and the animal was returned to the homecage. Tetrodes were lowered only once at each recording day.

A full-protocol experiment was composed of a sequence of six consecutive 10-min foraging sessions. The six sessions involved two standard light sessions in the presence of a proximal cue card (sessions “L1”-“L2”), a counterclockwise 90° cue rotation under standard light condition (session “R3”), another standard light session (session “L4”) followed by a dark session with cue removal (session “D5”) and a final standard light session (session “L6”). Between every two sessions (except before the dark session), animals were removed from the arena and allowed to rest for 5 min on the remote resting pot. During the dark session, however, while the animal was kept in the environment from the previous light session, the cue card was removed, the environment lights were turned off, and the recording began. The light and dark sessions were respectively carried out under 8-10 lux and 0.00 lux conditions (measured with a lux meter on the arena’s surface). Data analysis was performed offline. The automatically sorted spikes (Snap, Neuralynx Inc., USA) were refined manually using a graphical user interface software (Offline Sorter, Plexon Inc.) except for speed cells, for which, due to abundance, analysis was performed on stable automatically-sorted clusters. However, only clusters with an adequate isolation quality (Grade 2 or higher; Snap, Neuralynx Inc., USA). For each tetrode, the spike waveform features such as peak, valley, and height (peak-to-valley amplitudes), waveform energy, and principle components of spike waveforms were extracted and used to generate scatter-plots which was graphing a spike features of a channel against a spike features of another channel. These scatter plots provided visual aids to refine the automatically sorted clusters based on their waveform features. The quality of spike sorting for each isolated cluster was first tested by checking the absence of spikes in the refractory period (0-1ms). Further, to exclude poorly sorted clusters, an isolation distance measure was calculated for each cluster (Schmitzer-Torbert et al., 2005). Clusters with a low isolation distance score (below 10) were excluded from the analysis.

## QUANTIFICATION AND STATISTICAL ANALYSIS

### Construction of HD from samples

Two differently sized infrared LED circuits were designed and tested rigorously to ensure that the camera’s light detection makes them distinguishable in various behavioral or head rotation conditions (yaw, pitch, or during partial coverage by the arena boundary). However, the HD samples were further tested for a possible flip occurrence in the detection of LEDs in which bigger (rear) and smaller (front) LEDs were detected inversely. The lost HD samples were interpolated whenever the duration of sample loss was below 1.5 seconds and removed otherwise. The LED flip samples (when the big and small light spots were detected reversely) were detected and dealt similarly to the missing HD samples.

### HD rate-maps

The head direction sample was binned into sixty bins of 6°, and a “bin estimate” of HD rate map was determined as the number of spikes in each head direction bin divided by the amount of time spent in that bin (Taube, 1995). The rate map was then smoothed by a Gaussian kernel (σ = 7.2°).

### HD cell selection

HD cells were detected by multiple alternative methods, and a separate analysis was performed on cells classified by each of these methods indicating the same results. The methods include a generalized linear model framework for modeling spiking activity as a function of extrinsic covariates, including HD, and a shuffling procedure comparing the cell’s mean resultant-length or the Watson U^2^ value with a chance-level outcome driven from randomly shuffled spikes (each method is described in detail later). Only stable clusters that were detected in both two first standard sessions, “L1” and “L2” (stable clusters) and also identified as HD in both sessions (Stable HD firing), were classified as HD cells. Further, to exclude unstable or poorly recorded clusters, only cells with a minimum overall mean firing rate of 0.1 Hz in both two first sessions were included.

To ensure that the detected HD cell was not a byproduct of biased HD sampling from a cell with spatially selective firing, we further included the “distributive hypothesis” measure (Although not essential for HD cells selection by GLM; Cacucci et al., 2004; Kornienko et al., 2018; Muller et al., 1994). This method measures the similarity between the observed HD rate map and the predicted rate map from the HD occupancy probability over the spatial bins. The spatial scale of the environment was segmented into 2 × 2 cm spatial bins In (*i*^th^, *j*^th^) spatial bin, the firing rate of the cell facing direction is denominated as *R*_ij_(*ϕ*), and the time spent facing the direction *ϕ* in that spatial bin is denominated as *T*_ij_(*ϕ*). The predicted firing rate as a function of *ϕ* is defined as bellow:

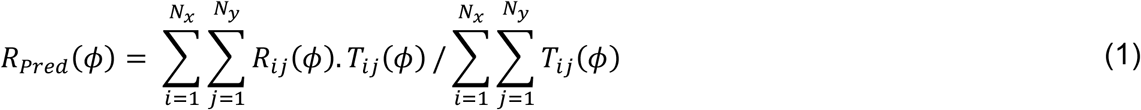

The distributive ratio was defined as:

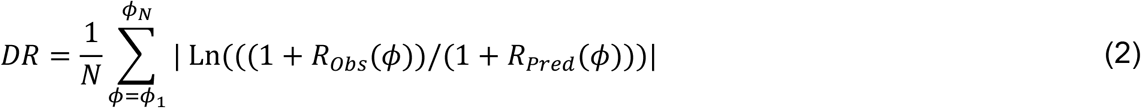

where *N* is the number of radial bins used for HD rate map (N = 60 for a bin size of 6°), *N*_x_ and *N*_y_ are the number of bins in spatial scale, and *R*_Obs_ is the bin estimate of HD rate map. A DR of zero indicates a complete prediction of the observed HD rate map by a combination of HD sampling and the spatial selectivity of cell firing. A high DR value, however, indicates that the head direction accounts for some variability of the rate map.

### Cell classification by shuffling

A shuffling test was performed to create a statistic for evaluating whether the cell firing can be significantly explained by an external covariate including speed, HD, and angular velocity (Kropff et al., 2015). The spike train of individual cells was time-shifted along with the tracking samples by a variable randomly chosen between 20 s and the total trial length minus 20 s. The shifted values that were passing the length of the trial were wrapped to the beginning. Such shuffling procedure decouples the spikes from external covariates (such as speed, HD, and angular velocity) while keeping the spike timing relations intact. For the shuffled spikes, a given parameter characterizing the influence of a given covariate was calculated. This procedure was repeated 100 times, and the distribution of the outcomes was generated. The 99^th^ percentile value of this shuffled distribution was used as the classification criterion. The parameters used for the classification of HD cells were either the mean vector length or Watson U^2^. For AHV and speed cells, a Pseudo-R^2^ value was measured after fitting a nonlinear curve fit (described in detail later) to the AHV and speed firing rate maps.

### Cell classifications by generalized liner model (GLM)

To quantify the effects and relative contribution of an extrinsic covariate (here, *i.e.*, head direction, angular head velocity, and linear speed) to the activity of a cell, while avoiding data sample binning and behavioral biases, the spiking activity was modeled as a discrete-time point process in the GLM framework under below intensity functions (Acharya et al., 2016; Kraus et al., 2015; Truccolo et al., 2005).

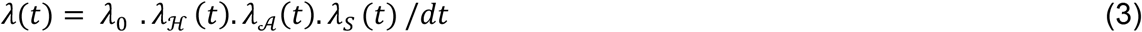

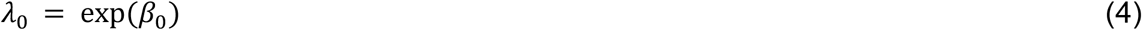

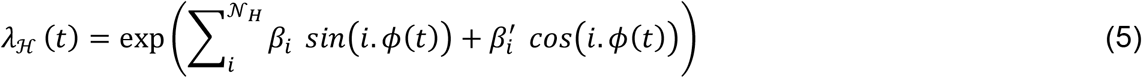

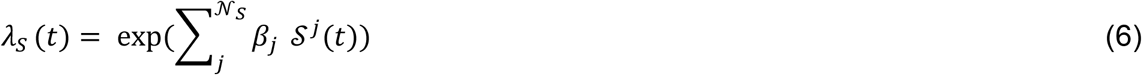

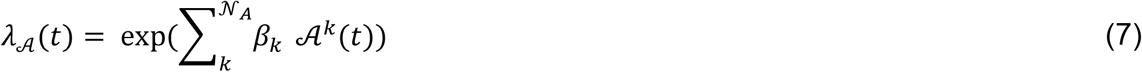

where *λ* stands for the firing rate of the cell. *λ*_0_, *λ*_ℋ_, *λ*_𝒜_, and *λ*_s_ stand for firing rates of baseline, head-direction, angular head velocity, and linear speed components in the model, respectively. *dt* is the time bin size (40 ms). *ϕ*(*t*), 𝒜(*t*) and 𝒮(*t*) are the instantaneous head direction, angular head velocity, and linear speed of the animal. 𝒩_ℋ_, 𝒩_𝒮_, 𝒩_𝒜_ are the model orders respectively for head-direction, angular head velocity, and linear speed components. *β*_0_, *β*_i_, *β*′_*i*_, *β*_j_, and *β*_k_ are the parameters of the model to be estimated.

Time bin size used to bin the spike trains was the same as the animal tracking sampling frequency (*dt*). To avoid under-sampled sections of data, AHV samples were limited to 1-99^th^ percentile and high-speed samples passing 99^th^ percentile of the speed sample distribution were also dismissed. Speed and AHV samples were scaled over the maximum bound within (0, 1) and (-1, 1), respectively. HD samples were converted to radian to bound within (0, 2*π*). The covariate matrix was built by including all covariate elements according to the model order and the intensity functions. Model fitting was performed in MATLAB using the function “glmfit” under a Poisson distribution assumption with a log link function. The maximum order numbers were preset at 𝒩_ℋ_ =4, 𝒩_𝒮_ =3, 𝒩_𝒜_ =3 and all combinations of models with orders of zero up to these maximum orders were tested.

To evaluate and choose the simplest model that has the best performance in estimating a cell’s spiking, we used the Bayesian Information Criterion (BIC), an information-based criterion that penalizes models with more parameters (Acharya et al., 2016; LaChance et al., 2019; Wang et al., 2018). Note that, for instance, in a one-component H model with an order of four, the number of terms is higher than the same model with one order, and likelihood alone would not penalize this complexity.

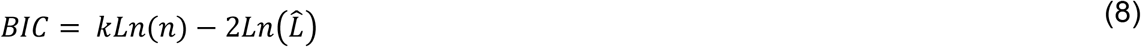

Where *k* is the number of parameters estimated by model (total number of *β* parameters), *L̂* is the maximized value of the likelihood function and *n* is the number of the sample size. Further, to avoid overfitting by including more components for each cell, we performed a ten times ten-fold cross-validation procedure by dividing the data samples into the train and test sets (Fallahnezhad et al., 2011; Hardcastle et al., 2017). Models were fit on train sets, and comparisons for model selection were made over test samples. We performed a nested method to select the best model starting from the simplest model (an intercept model) and checking if any model with an extra component can perform better. Therefore, we first fitted all the models with one component (*i.e., four* ℋ models, three 𝒮 models and three 𝒜 models). Among all these models, the one with the lowest average BIC value (over ten sections of test data samples) was chosen as the best model with one component and was compared to an intercept model using a one-sided signed-rank test with an a level of 0.05. If this model was were was significantly better than an intercept model, then the procedure would continue by comparing the best one-component model with models containing two components (ℋ𝒮, ℋ𝒜, 𝒮𝒜) but only those containing the best one-component model. For instance, if the best one-component model was ℋ, then the best model among all models (means a combination of all different order sizes for each component) forming either of ℋ𝒮 or ℋ𝒜 would be compared to the ℋ-model..

Finally, a “GLM estimate” of rate map for a cell was reconstructed as below:

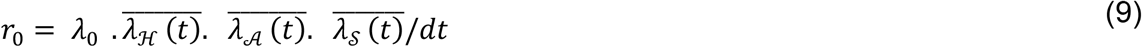

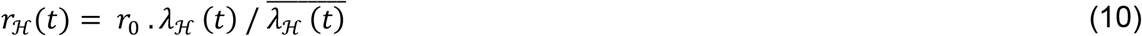

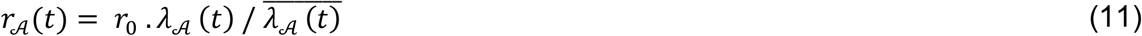

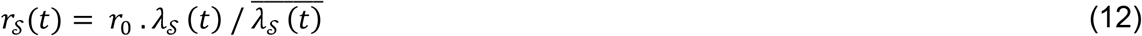

Where *r*_0_ is the baseline firing rate, *r*_ℋ_, *r*_𝒜_, and *r*_𝒮_ are the reconstructed rate maps associated with HD, AHV, and speed components, respectively, and 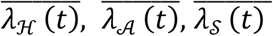 are the mean values of reconstructed intensity functions. The GLM estimate of HD, AHV, and speed rate-maps were constructed by calculating intensity functions (formulas 4-7) and firing rate functions (formulas 9-12) by setting values within (0, 2*π*), (0, 1), and (-1, 1) to the input variables *ϕ*(*t*), 𝒮(*t*), and 𝒜(*t*), respectively.

### HD rate-map properties

*Mean* v*ector length:* the magnitude of the resultant vector obtained from the head direction angles collected during the spiking of a cell.

*Preferred firing direction (PFD)*: the angle of the resultant vector obtained from the head direction angles during the spiking of a cell.

*Intra-session stability of an HD rate-map:* Sessions were divided into two 5-min (Figures 1 and 2) or five 2-min sub-sessions (Figure S1). Stability as a correlation measure was evaluated by Pearson correlation between HD rate maps of these sub-sessions (for five 2-min sub-sessions, an average of Pearson correlation between either all pairs or consecutive pairs was calculated). The angular variance between PFDs of 2-min sub-sessions was also calculated, representing a reciprocal measure to the stability (Figure S1).

*Inter-session stability of an HD rate-map:* Pearson correlation value between HD rate maps of two sessions.

*Burst index:* the percentage of spikes occurring within an interval smaller than 6 ms to total spike count.

*Theta index*: extracted from FFT-based power spectral density of spike-train autocorrelation signal (Langston et al., 2010). Spike-time autocorrelation for individual cells was generated from spike trains binned in 2 ms with lags up to 2 s. The value on zero lag was set to zero. The normalized distribution of autocorrelogram was generated by subtracting the mean value from all values divided by the sum of all values. AThe autocorrelation signal was tapered with a Hanning window to reduce spectral leakage. The FFT was scaled to the length of the autocorrelation signal. The power spectrum was generated by FFT in MATLAB. The power spectrum was set as the square magnitude of the FFT signal. Theta index was defined as the ratio of the average power spectrum in the theta frequency band (4-11 Hz) to the average spectrum in frequencies between 0 Hz to 125 Hz. Cells with theta index greater than 5 were considered theta modulated.

*Information (bits/spike):* Analogous to information entropy, head-direction information was defined as (Skaggs et al., 1993):

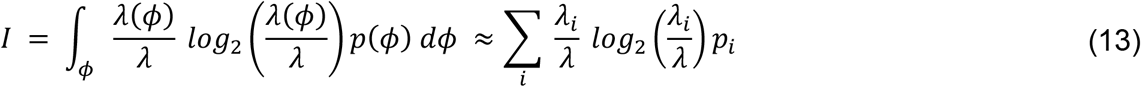

where *I* is the information rate of the cell in bits per spike, *ϕ* is the head direction, *p*(*ϕ*) is the probability density of head pointing toward direction *ϕ*, *λ*(*ϕ*) is the mean firing rate when the mice head is pointing toward the direction *ϕ*, *i* is the radial bin index (*i*-th bin of *ϕ*), *p*_i_ is occupancy probability in bin *i*, *λ*_i_ is mean firing rate in bin *i*, and *λ* is overall mean firing rate.

### AHV and Speed cells

As a first approach, the GLM framework was used to classify AHV and Speed cells along with HD cells. As a second approach to generate a Speed or AHV score, we modeled the firing rate of the cells with a single-component (speed or AHV) input using the maximum likelihood estimator (Hinman et al., 2016; Sharp et al., 2001; Stackman and Taube, 1998). We assumed the firing rate of a speed cell follows either a linear or a saturating exponential function (Hinman et al., 2016), and for an AHV cell, either a linear (symmetric AHV), parabola, or hyperbola (asymmetric AHV) function. The fitted functions were optimized using the “mle” function in MATLAB and custom-made code adopted from the codes available online (Hinman et al., 2016). F-statistic and p-values for each fit were calculated from log-likelihood against an intercept model. These measures were also computed for fitted models against each other to find the model with the best fit. For a speed cell, the function that fits better than both an intercept model and the alternative model was chosen. For AHV cells, a nested comparison starting from the simplest model to a more complex model was made. Specifically, starting from the linear model, if the model was not performing better than the intercept model, the intercept model was chosen (no AHV component). If the linear model was performing better than an intercept model, the comparison was continued by fitting a parabola and comparing the fit with that of the linear model, and so on. A Pseudo-R^2^ value based on Nagelkerke/Cragg & Uhler’s method was calculated to assess the goodness of a fit:

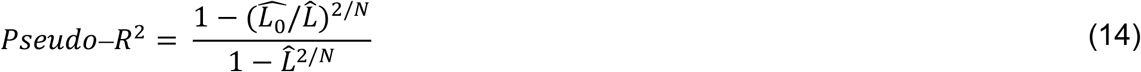

where *N* is the number of samples and 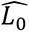 and *L̂* are the likelihood of the intercept model and fitted model, respectively.

To evaluate whether the speed or AHV component significantly accounts for some variability in observed spiking of the cell, we assessed whether the fitted model performs significantly better than a chance-level statistic. A spike shuffling procedure created the chance-level statistic described earlier. The selected model was then compared to models with the same orders fitted to the shuffled data, and the 99^th^ percentile of Pseudo-R^2^ values of shuffled data was used as the criterion.

Finally, we included an additional criterion to limit speed and AHV cell comparisons only to the structures where HD cells are recorded. For speed cells of the ADN group, due to abundance and sufficient sample size, we limited the speed cell samples to the clusters which were co-recorded with at least one cluster that was identified as HD cell in at least one of the first two standard light sessions. For AHV cells of the ADN group, we used the same criteria as speed cells. However, we also included extra AHV cells recorded in between experimental days, where at least one HD cell was recorded in both days. No similar criteria were set for the speed and AHV cell samples of RSC groups as far as post hoc histological analysis confirmed the tetrode path within the structure.

### Temporal coordination between HD cells by spike cross-correlogram

A deterministic correlation for a cell pair (two simultaneously recorded cells) was generated from spike trains binned in 100ms or 200ms with lags up to 20s using “xcorr” in MATLAB (Bassett et al., 2018; Peyrache et al., 2015). To create a comparable situation between different pairs and remove the effect of various firing rates, which can lead to variable peaks in the histogram, cross-correlograms were normalized by a chance-level cross-correlogram (Bassett et al., 2018). In this regard, the spike train of one of the cells in a pair was time-shifted by a variable randomly chosen between 20 s and the total trial length minus 20 s. The shifted spikes that were passing the original size of the trial were wrapped to the beginning. Similar to original spike trains, a cross-correlogram was calculated for the shifted spikes but with lags of ±1 s. This procedure was repeated 500 times, and the average of the 500 cross-correlograms was considered a chance-level distribution for the central part (±1 s) of the cross-correlogram. The normalized cross-correlogram was generated by division of the original cross-correlogram from actual spikes to the chance-level distribution. Further, the mean value of the central 1-second of the normalized cross-correlogram was extracted. This was referred to as *m_xco_* and is used to examine the stability of the HD network correlation structure between different sessions.

### Temporal coordination between HD cells by Kendal’s correlation

Since a cell’s firing is typically sparse when performing cross-correlation over the spike train of a pair of cells in a short time window, the co-firing histogram can contain a large number of zero-zero associations for the pair’s co-firings. This will create a prominent association for the pair’s firing and can mask the accurate degree of co-firing and create a significant correlation value when using a parametric measure. The non-parametric Kendall’s correlation, however, incorporates an adjustment for ties with a normalization constant (tau-b). Spike trains were first binned in either of 250 or 1000 ms, and Kendall’s correlation coefficient (*τ_Kendal_*) was measured for a simultaneously recorded cell pair (only the results for 1000 ms binning window were reported, albeit both binning yielded the same results). *τ_Kendal_* is used to examine the stability of the HD network correlation structure between different sessions (Park et al., 2019).

### Stability of temporal coordination in HD cell network

To compare the correlation structure existing between HD cells from one session to another, the correlation value (either *m_xco_ or τ_Kendall_*) for all cell pairs from one session was plotted against another session as a scatter plot. The stability of correlations between two sessions was quantified by Pearson correlation (*ρ*) between the correlation values (*m_xco_* or *τ_Kendall_* of cells pairs) of those two sessions (Park et al., 2019). Further, to compare the stability of such correlation structure between multiple sessions, the distribution of Pearson correlation between session L2 and L1 (*ρ*_*L*1X*L*2_) was used as observed baseline stability of the correlated HD network and the distribution of Pearson correlations between any other session with session L1 was compared to this baseline (e.g. *ρ*_*L*1X*L*4_ *or ρ*_*L*1X*D*S_).

### Statistics

Multiple factor comparisons were performed by ANOVA, under a mixed-design model assumption (IBM SPSS). Statistical comparisons between distribution of correlations (r) were performed by a custom-made MATLAB program as two-tailed comparisons on Fisher z-transformed correlation values ( z = 0.5 ln((1+r)/(1-r)) ). Circular statistics were performed with the CircStat Matlab toolbox (Berens, 2009). Permutation tests for creating chance-level statistics were performed by custom-made programs in MATLAB. The significance level was set to 0.05.

### Histological analysis

After the completion of experiments, animals were deeply anesthetized with an overdose of sodium pentobarbital and perfused intracardially with saline and 4% paraformaldehyde (PFA) in a 0.1M phosphate buffer. To keep tetrode tracks intact, the tetrodes were first brought up out of the brain by turning the screws on the micro-drives, and then the brain was extracted. The extracted brain was stored in PFA overnight at 4°C and then cryoprotected in a series of sucrose solutions (10%, 20%, and 30%, all in phosphate-buffered saline) for a minimum of 3 days at 4°C. Brains were frozen and fixed on a cryostat, and coronal slices of 40-µm thick were cut from both ADN and RSC structures and collected on gelatin-coated slides. Collected slices were stained with cresyl violet solution (Sigma) and protected by a mounting medium (Eukitt, Sigma) and coverslips. Brightfield images were taken with a microscope (Leica DM 5500 B) equipped with 2.5X and 5X objectives.

